# Clonal dynamics of SARS-CoV-2-specific T cells in children and adults with COVID-19

**DOI:** 10.1101/2022.01.30.478400

**Authors:** Weng Hua Khoo, Katherine Jackson, Chansavath Phetsouphanh, John J. Zaunders, José Alquicira-Hernandez, Seyhan Yazar, Stephanie Ruiz-Diaz, Mandeep Singh, Rama Dhenni, Wunna Kyaw, Fiona Tea, Vera Merheb, Fiona X. Z. Lee, Rebecca Burrell, Annaleise Howard-Jones, Archana Koirala, Li Zhou, Aysen Yuksel, Daniel R. Catchpoole, Catherine L. Lai, Tennille L. Vitagliano, Romain Rouet, Daniel Christ, Benjamin Tang, Nicholas P. West, Shane George, John Gerrard, Peter I. Croucher, Anthony D. Kelleher, Christopher G. Goodnow, Jonathan D. Sprent, Joseph D. Powell, Fabienne Brilot, Ralph Nanan, Peter S. Hsu, Elissa K. Deenick, Philip N. Britton, Tri Giang Phan

## Abstract

Children infected with severe acute respiratory syndrome coronavirus 2 (SARS-CoV-2) develop less severe coronavirus disease 2019 (COVID-19) than adults. The mechanisms for the age-specific differences and the implications for infection-induced immunity are beginning to be uncovered. We show by longitudinal multimodal analysis that SARS-CoV-2 leaves a small footprint in the circulating T cell compartment in children with mild/asymptomatic COVID-19 compared to adult household contacts with the same disease severity who had more evidence of systemic T cell interferon activation, cytotoxicity and exhaustion. Children harbored diverse polyclonal SARS-CoV- 2-specific naïve T cells whereas adults harbored clonally expanded SARS-CoV-2-specific memory T cells. More naïve interferon-activated CD4^+^ T cells were recruited into the memory compartment and recovery was associated with the development of robust CD4^+^ memory T cell responses in adults but not children. These data suggest that rapid clearance of SARS-CoV-2 in children may compromise their cellular immunity and ability to resist reinfection.

**HIGHLIGHTS:**

- Children have diverse polyclonal SARS-CoV-2-specific naïve T cells
- Adults have clonally expanded exhausted SARS-CoV-2-specific memory T cells
- Interferon-activated naïve T cells differentiate into memory T cells in adults but not children
- Adults but not children develop robust memory T cell responses to SARS-CoV-2

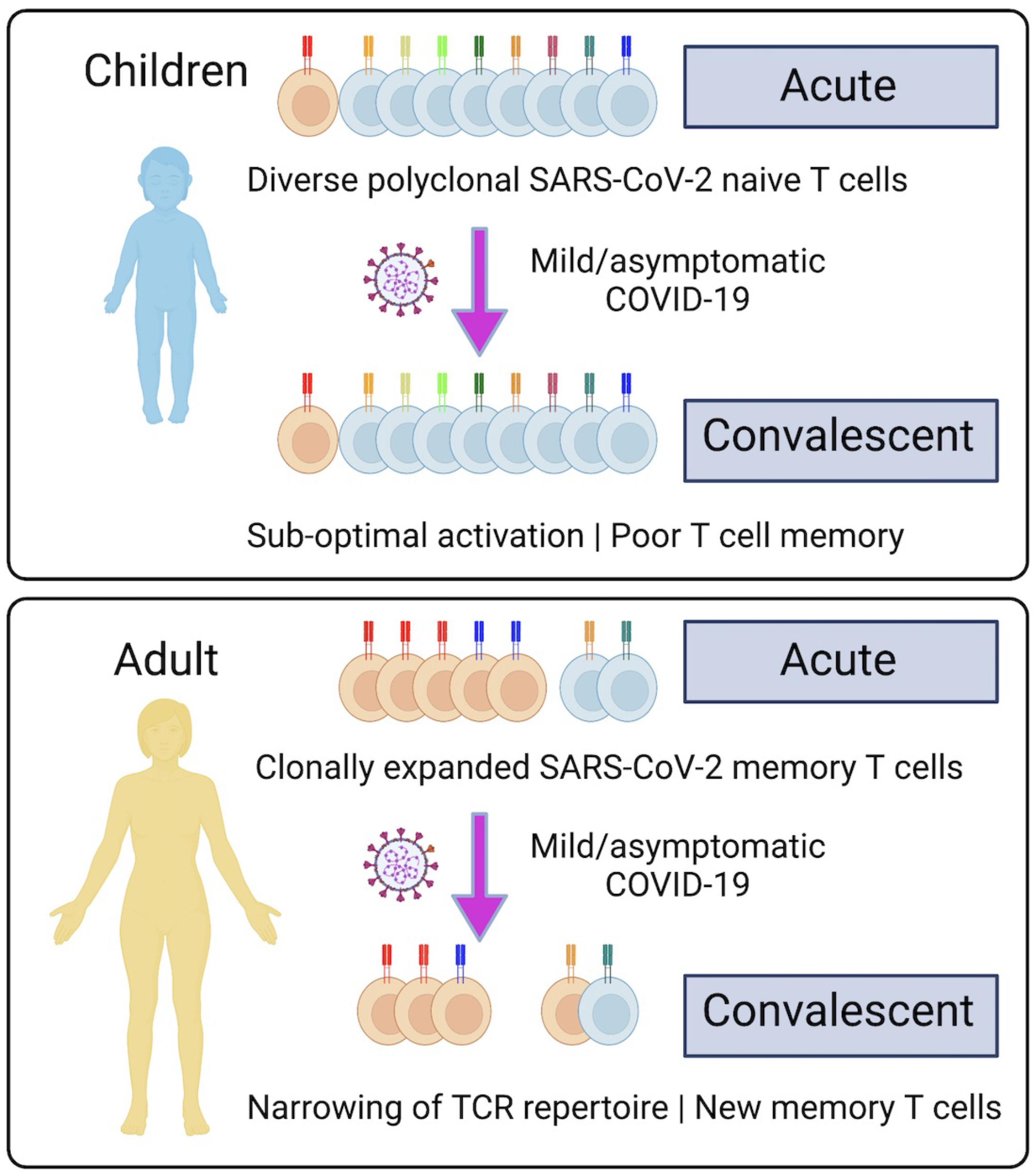

## Introduction

Infection with the respiratory pathogen severe acute respiratory syndrome coronavirus 2 (SARS- CoV-2) causes the pandemic coronavirus disease 2019 (COVID-19) (Guan et al., 2020; Zhou et al., 2020). Disease severity varies widely from asymptomatic infection in the majority of individuals, to severe life-threatening disease in a minority of patients (Chen et al., 2020; Huang et al., 2020; Wang et al., 2020). Older age, male sex and comorbidities such as hypertension, cardiovascular disease and diabetes have been identified as independent risk factors for severe disease and death (Jordan et al., 2020). There is now consistent evidence across multiple different settings and locations that the severity of COVID-19 infection increases substantially with age (O’Driscoll et al., 2021).

Recently, a number of investigators have examined the local and systemic immune responses of children to SARS-CoV-2 to determine the potential mechanisms for these age-specific differences in susceptibility to infection and disease. Most notably, it has been shown that children have an enhanced antiviral sensing and stronger antiviral interferon response in the upper airways due to higher basal expression of the MDA5 and RIG-1 viral pattern recognition receptors and interferon gene signatures in nasal epithelial cells, macrophages and dendritic cells (Loske et al., 2021; Yoshida et al., 2021). This pre-activated innate immune system may be more efficient at clearing SARS-CoV-2 infection. Indeed, analysis of three child household contacts of adults with PCR-confirmed symptomatic COVID-19 suggested that children may mount an effective early antiviral immune response that eliminates the virus without any detected PCR evidence of SARS-CoV-2 infection (Tosif et al., 2020). These differences in the innate immune response of children may also be detectable in the immunophenotype of circulating neutrophils, dendritic cells, monocytes and natural killer (NK) cells in the peripheral blood (Neeland et al., 2021).

The adaptive immune response to SARS-CoV-2, particularly the humoral component provided by antibodies and memory B cells, can be either protective or pathogenic in COVID-19 (Bartsch et al., 2021; Zohar and Alter, 2020). There is evidence that antibodies against endemic human coronaviruses (hCoV) cross-react with SARS-CoV-2 and these may be back-boosted upon infection (Anderson et al., 2021; Aydillo et al., 2021; Ng et al., 2020). However, this pre-existing cross-reactive humoral immunity does not appear to provide protection, and may potentially be harmful by locking in the memory B cell responses and preventing the emergence of *de novo* naïve B cell responses to novel SARS-CoV-2 antigens, a phenomenon known as original antigenic sin (Aydillo et al., 2021; Dhenni and Phan, 2020; Francis, 1960; Zhang et al., 2019). Analysis of pre-pandemic serum has also shown that healthy elderly individuals have higher immunoglobulin class-switched IgA and IgG antibodies that cross-react with SARS-CoV-2, whereas children have elevated cross-reactive SARS-CoV-2 IgM antibodies (Selva et al., 2021), suggesting that they have less exposure to hCoV and are less antigen- experienced but more polyreactive. Furthermore, it has been suggested that pathogenic responses associated with severe COVID-19 are linked to SARS-CoV-2 IgA and neutrophil hyperactivation (Bartsch et al., 2021) and that the antibodies produced by children differ from adults in their Fc- dependent antibody effector functions such as antibody-dependent cellular cytotoxicity, phagocytosis and complement activation (Bartsch et al., 2021; Selva et al., 2021).

There is emerging evidence that cellular immunity against SARS-CoV-2 provided by T cells may be similarly impacted by prior exposure to hCoV. Using overlapping peptide pools for *in vitro* T cell stimulation assays, investigators have detected CD4^+^ and CD8^+^ T cells cross-reactive against SARS- CoV-2 spike (S), membrane (M), nucleocapsid (N) and open reading frames (ORF) in pre-pandemic blood samples from unexposed individuals (Bacher et al., 2020; Braun et al., 2020; Grifoni et al., 2020; Le Bert et al., 2020; Mateus et al., 2020). Consistent with this, SARS-CoV-2 T cell responses have been shown to be lower in children and increase with age and time after infection (Cohen et al., 2021).

We performed longitudinal analysis of the immune response of seven children and five adults from the same household with mild/asymptomatic COVID-19 in the community confirmed by reverse transcription polymerase chain reaction (RT-PCR) and an additional two unrelated adults who were ventilated in the intensive care unit with severe life-threatening COVID-19. We analyzed the cellular phenotype, serum antibody response to SARS-CoV-2, cytokine profile, *in vitro* memory T cell responses to recombinant S and RBD proteins, and simultaneous single cell transcriptome and TCR and B cell receptor (BCR) repertoire sequencing of 433,301 single cells obtained from acute and convalescent blood samples. These multimodal analyses identified novel subpopulations of naïve T cells, including acutely expanded clusters of interferon-activated naïve T cells which differentiate into memory T cells in convalescence. We show that mild/asymptomatic COVID-19 results in systemic activation of both the innate and adaptive immune compartments in adults. In contrast, children had less activation of the circulating T and B cells. Children have more SARS-CoV-2- specific T cells which predominantly have a naïve phenotype and diverse TCR repertoire. Adults have fewer SARS-CoV-2-specific T cells which are more antigen-experienced and often harbor clonally expanded exhausted memory T cells. Circulating T cells in children retain their naïve state and did not generate many antigen-specific memory T cells despite infection with the virus. In contrast, adults generated more memory T cells from the naïve interferon-activated and this was associated with the development robust SARS-CoV-2-specific memory T cell responses in adults, but not children. This failure of infection-induced immunity places children at the risk of recurrent infection and progressive restriction of their T cell repertoire and responses as they grow older.

## RESULTS

### Longitudinal tracking of the immune response to SARS-CoV-2 in children and adults

Acute and convalescent blood samples from seven children (<16 years of age) and five adults (>30 years of age) with mild/asymptomatic disease (WHO Clinical Progress Scale of 0 or 1 out 10), and two adults who were intubated and ventilated in the intensive care unit (ICU) with severe disease (WHO Clinical Progress Scale of 7 out of 10) were analyzed (**Fig. 1A, 1B**). Two of the children (C3 and C4) were identical twins. Both children and adults developed antibodies to S protein in the convalescent phase, with the highest antibody titres detected in the ICU adults (**Fig. 1C**). High dimensional flow cytometry showed consistent differences in the distribution of circulating natural killer (NK) cells, naïve and memory B and T cells that reflected age-specific differences between children and adults (**Fig. S1**). Nevertheless, there was evidence of increased T cell activation in the acute stage with 2-fold expansion of activated CD38^+^HLA-DR^+^ CD4^+^ T cells and 3-fold expansion of CD38^+^HLA-DR^+^ CD8^+^ T cells in non-ICU adults compared to children (**Fig. 1D**). ICU adults had the highest frequency of CD38^+^HLA-DR^+^ T cells. Serum cytokine analysis detected 50- to 100-fold elevation of interleukin-6 (IL-6) in the ICU adults compared to acute children, convalescent children, acute non-ICU adult and convalescent non-ICU adult (p=0.001, one-way ANOVA with Tukey’s corrections) (**Fig. 1E**).

**Figure 1.**
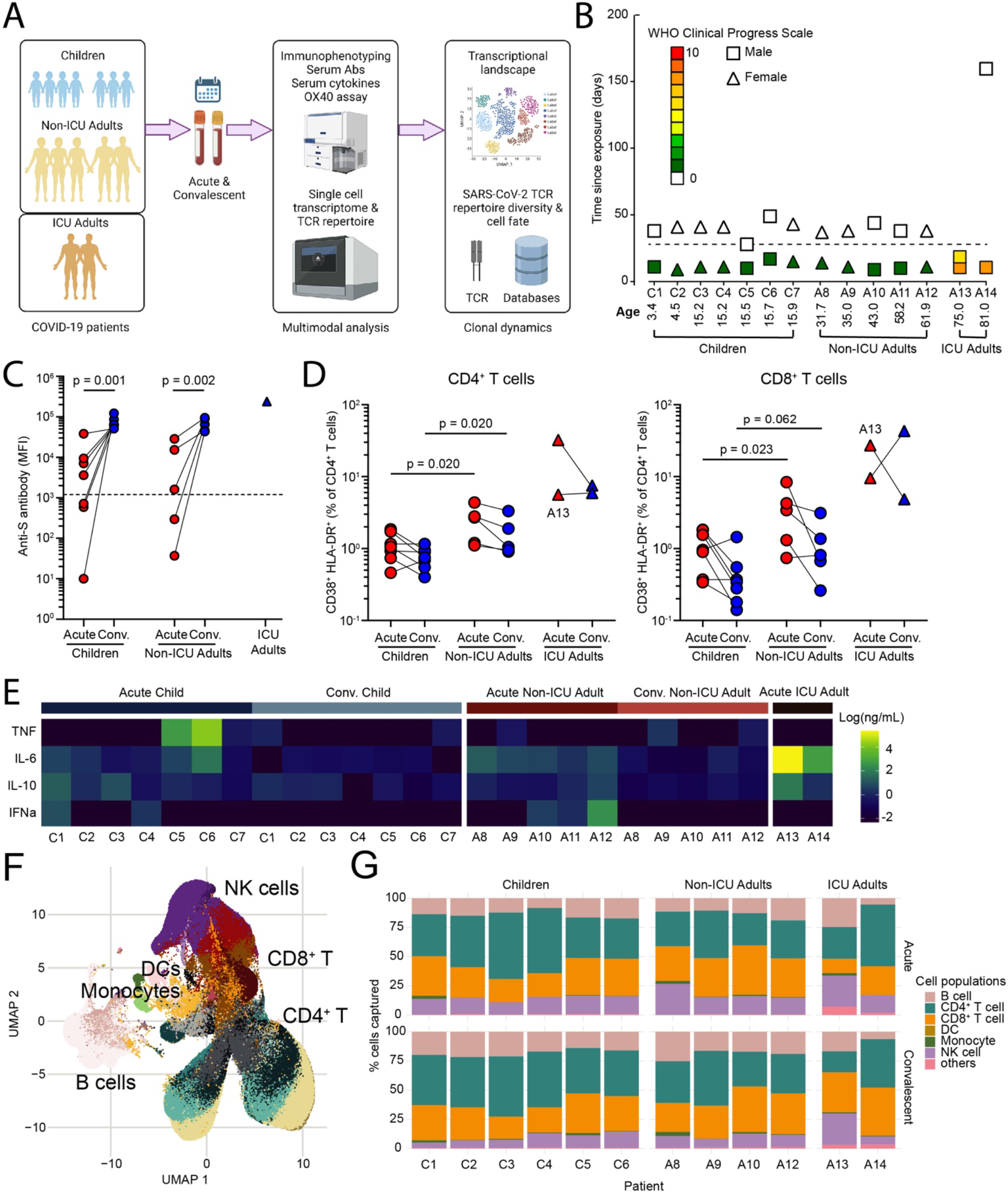
Multimodal analysis of children and adults with mild COVID-19. (A) Study overview. (B) Patient demographic and clinical severity score. (C) Serum anti-S protein antibodies. (D) CD38^+^HLA-DR^+^ activated CD4^+^ (left) and CD8^+^ (right) T cells in children, non-ICU and ICU adults during Acute and Convalescent phases. (E) Heatmap of serum TNF, IL-6, IL-10 and interferon-*α*. (F) UMAP showing 433,301 single cells from children, non-ICU and ICU adults during Acute and Convalescent phases. (G) Stacked barplot showing cellular composition of PBMCs in children, non-ICU and ICU adults during Acute (top) and Convalescent (bottom) phases.

We next generated single cell transcriptomes from acute and convalescent samples from these 14 COVID-19 patients. In total, we sequenced 522,926 cells and excluded samples from two patients (C7 and A11 from the same household) due to batch effects (**Fig. S1C**). The remaining 433,301 cells were visualized in 2D Euclidean space by uniform manifold approximation and projection (UMAP) (**Fig. 1F**). We initially annotated clusters of cells which consisted of populations of 69,074 B lineage cells, 171,393 CD4^+^ T cells, 127,069 CD8^+^ T cells, 376 dendritic cells (DCs), 4,135 monocytes, 56,807 NK cells, and 4,447 other cell types (including HSCs, platelets, erythrocytes and doublets) (**Fig. 1G** and **Fig. S2**). Analysis of the cellular composition identified similar age-specific changes to the circulating NK, B and T cell populations as observed by the flow cytometry (**Fig. 1G** and **Table S1**). Taken together, these data show that, in addition to age-specific differences in the circulating immune compartments in children and adults, there is evidence for greater systemic T cell activation in adults.

### Deconvolution of the circulating immune compartment in children and adults with COVID-19

We resolved the circulating innate and adaptive immune compartments in children and adults into populations of DCs (plasmacytoid dendritic cells, pDC; AXL^+^SIGLEC6^+^ dendritic cells, ASDCs; type 2 conventional dendritic cell, cDC2), NK cells (proliferating; CD56^dim^; CD56^bright^), monocytes (CD14^+^; CD16^+^), other cells (platelets; erythroblasts; HSPCs; innate lymphoid cells, ILCs; doublets), B lineage cells (naïve, memory, intermediate and plasmablasts), CD4^+^ T cells (naïve; stressed naïve; transcriptionally active naïve; interferon-activated naïve; CD40LG^+^ naïve; early memory; effector memory (TEM); early memory and regulatory T cells (Treg), and CD8^+^ T cells (naïve; PECAM-1^+^ naïve; interferon-activated naïve; TEM; GZMK^+^ TEM; CD20^+^ TEM; cytotoxic; KLRB1^+^ cytotoxic T cells) (**Fig. 2A**). Semi-automated annotation of the T cell compartment based on the expression of canonical genes revealed marked heterogeneity, particularly in the naïve T cell compartment with several previously unappreciated subpopulations of naïve CD4^+^ and CD8^+^ T cells that showed evidence of TCR-independent bystander activation by upregulation of CD40LG, stress-induced genes, transcriptional activity and interferon-induced genes (**Fig. 2B**). Analysis of the genes expressed by the different T cell clusters revealed interferon-activated naïve CD4^+^ and CD8^+^ T cells express a number of interferon-induced genes including *IFI44L, MX1, ISG15, IRF7* and *OAS1* (**Fig. 2B** and **Table S2**). Similarly, we identified a number of memory and effector subpopulations including CD8^+^ GZMK^+^ TEM and KLRB1^+^ cytotoxic cells based on their transcriptional profile and TCR receptor usage. CD8^+^ GZMK^+^ TEM are a newly described subset of exhausted-like memory T cells expressing *TOX, TIGIT, FGFBP2, TBX21, SLAMF7, ZEB2, NKG7, GZMH* and *PRF1* (**Table S2**) that give rise to dysfunctional, exhausted effector T cells (Galletti et al., 2020). KLRB1^+^ cytotoxic T cells include MR1-restricted mucosal-associated invariant T (MAIT) cells expressing TRAV1/TRAJ33 and IL-17-producing cytotoxic T cells (Tc17) expressing *MAF, RORC, TBX21, EOMES, IL18R1, CCR6, PRF1, GZMK, GZMA, NKG7, CST7* and *GNLY* (**Table S2**). KLRB1^+^ cytotoxic T cells are reported to be tissue-homing IL-17A producing cells (Billerbeck et al., 2010); however, we did not detect upregulated expression of *IL17A* or related IL-17 family genes.

**Figure 2.**
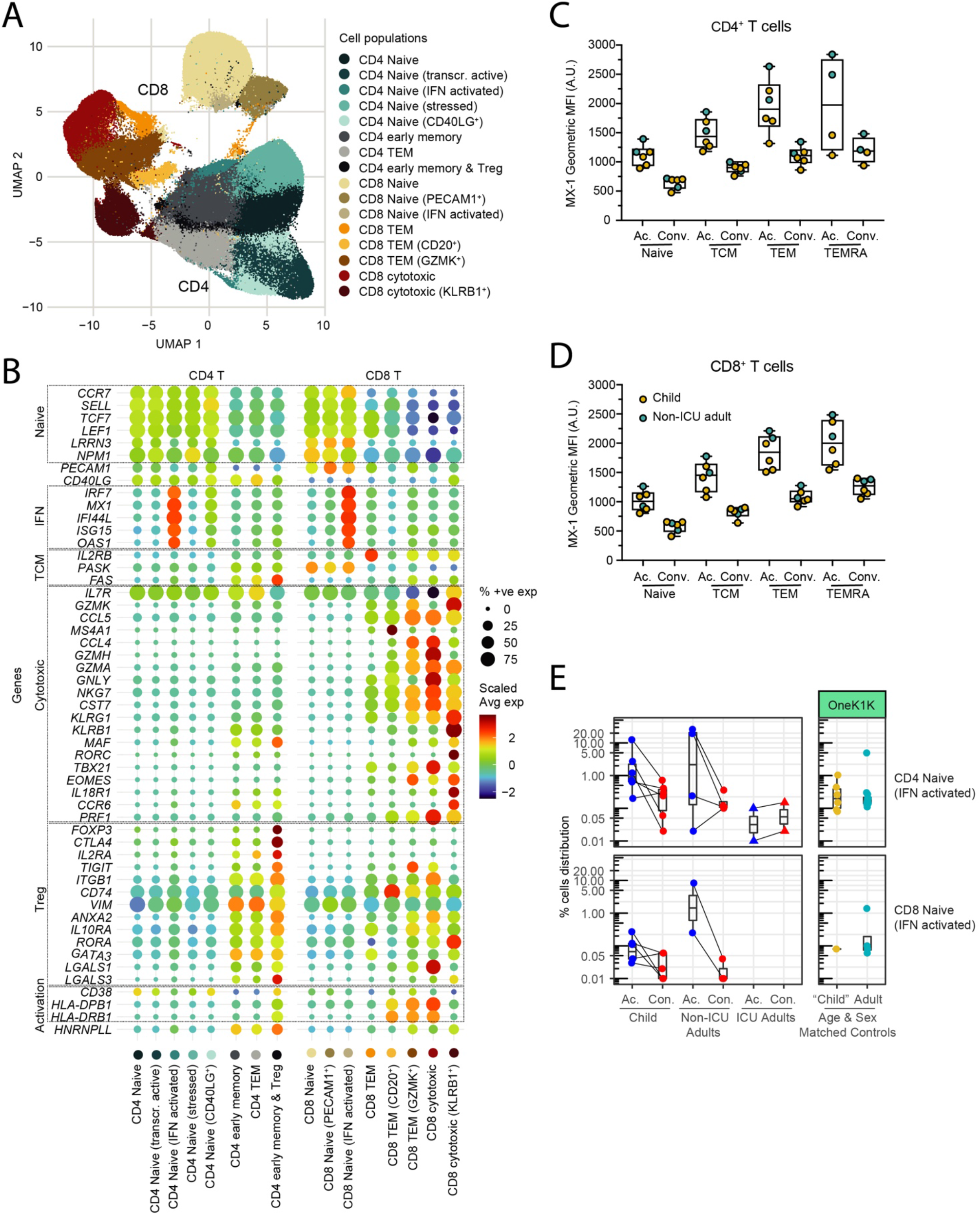
Decomposition of the T cell compartment in children and adults with COVID-19. (A) UMAP showing 171,393 CD4^+^ T cells and 127,069 CD8^+^ T cells from children, non-ICU and ICU adults during Acute and Convalescent phases. (B) Expression of genes associated with the naïve, interferon response (IFN), T central memory (TCM), T cell cytotoxicity, regulatory T cell (Treg) and activation states by different subclusters of CD4^+^ and CD8^+^ T cells. (C) Detection of interferon-induced MX-1 protein in subpopulations of CD4^+^ T cells during Acute and Convalescence. Naïve = CD45RA^+^CD45RO^−^; TCM = CD45RA^−^CD45RO^+^CCR7^+^CD62L^+^; TEM = CD45RA^−^CD45RO^+^CCR7^−^CD62L^−^; TEMRA = CD45RA^+^CD45RO^−^CCR7^−^CD62L^−^. (D) Detection of interferon-induced MX-1 protein in subpopulations of CD8^+^ T cells during Acute and Convalescence. T cell markers are as in (C). (E) Detection of interferon-activated naïve CD4^+^ T cells (top) and CD8^+^ T cells (bottom) in children, non-ICU and ICU adults during Acute and Convalescent phases (left) and in healthy age and sex- matched donors in the OneK1K cohort (right).

Intracellular flow cytometry confirmed the upregulated expression of the interferon-inducible MX-1 protein in a number of cell types, including naïve CD4^+^ (**Fig. 2C**) and CD8^+^ T cells (**Fig. 2D**), particularly in the acute phase. To determine if these subpopulations of naïve T cells were unique to COVID-19 or were also present in individuals who had not been exposed to SARS-CoV-2, we examined the distribution of T cells in age-matched PBMCs from the OneK1K cohort which was collected pre-2019 (**Fig. 2E**). This analysis showed that these novel interferon-activated naïve CD4^+^ and CD8^+^ T cells are also present in healthy adults who have not been infected with SARS-CoV-2. Thus, the circulating T cell compartment in children and adults is heterogeneous with multiple previously unrecognized subpopulations of naïve and activated T cells, including interferon-activated naïve CD4^+^ and CD8^+^ T cells.

### Differential gene expression between children and adults infected with SARS-CoV-2

We analyzed for differentially expressed genes (DEGs) between each subpopulation to determine the differences between children and non-ICU adults who both had mild/asymptomatic COVID-19. We detected very few DEGs in the adaptive B and T cell compartments in children compared to adults as they transitioned from acute to convalescence (6 upregulated, no downregulated genes in children; 22 upregulated, 43 downregulated genes in non-ICU adults; p=0.003, Fisher’s exact test) (**Fig. 3A**). In contrast, there were a large number of DEGs in the innate NK cell and monocyte compartment in both groups (25 upregulated, 24 downregulated genes in children; 67 upregulated, 50 downregulated genes in non-ICU adults; p=0.285, Fisher’s exact test). Notably, there was upregulation of interferon- induced genes (e.g. *IFI66, IFI44L* and *XAF1*) during acute infection and mitochondrial oxidative phosphorylation (OXPHOS) genes (e.g. *MT-ATP6, MT-CYB, MT-ND4* and *MT-CO1*) upon recovery (**Fig. S2**). In contrast, there were a large number of differentially expressed genes in both the innate and adaptive immune compartments in non-ICU adults (**Fig. 3A**). Notably, there was significant upregulation of interferon-induced genes in the B, CD4^+^ and CD8^+^ T cell compartments, as well as the NK cell and monocyte compartments, during acute infection, and upregulation of OXPHOS genes upon recovery (**Fig. S2**). Direct comparison of acute children and acute non-ICU adult PBMCs revealed the largest differences were in the monocyte and CD8^+^ T cell compartment (**Fig. 3A** and **Fig. S2**). There was upregulation of naïve/T memory stem cell genes (e.g. *IL7R, LEF1, NOSIP, SELL* and *TCF7*) in CD8^+^ T cells in children and upregulation of cytotoxicity genes (e.g. *CST7, GNLY, GZMA, GZMB, GZMH, NKG7* and *PRF1*) in non-ICU adults. These acute gene expression differences between children and non-ICU adults in the CD8^+^ T cell compartment were less prominent but nevertheless persisted into convalescence (**Fig. S2**). We next performed DGE analysis on each cell subpopulation (**Fig. 3B**). There was a significant number of differentially expressed genes for all cell clusters except for the DC subsets and the interferon-activated naïve CD8^+^ T cells. Importantly, greater gene expression differences were observed in non-ICU adults than in children, particularly in the NK, B and CD8^+^ T cell subpopulations (**Fig. 3B** and **Fig. S2**).

**Figure 3.**
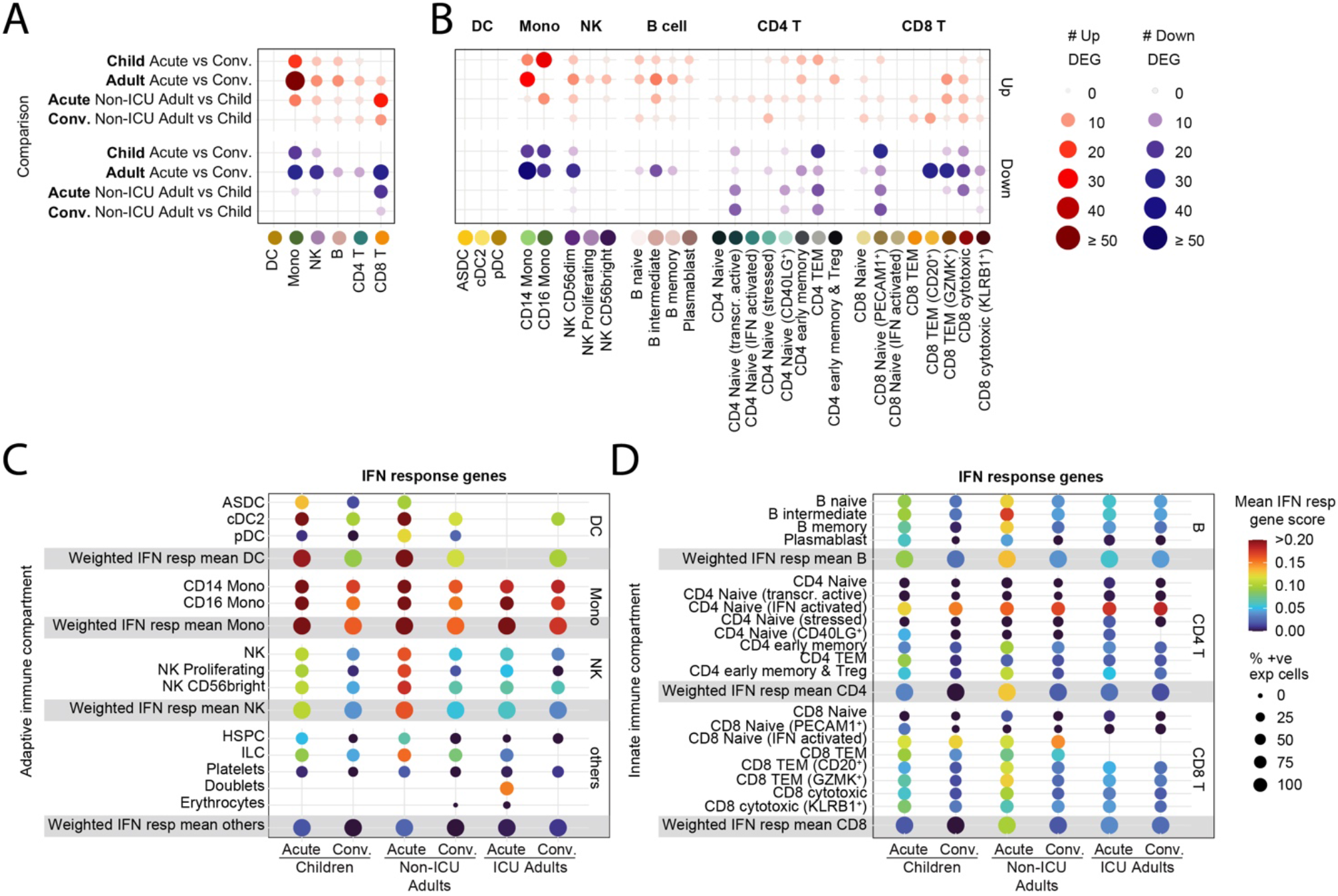
Transcriptomic differences between children and adults with mild COVID-19. (A) Dotplot showing the number of differentially expressed genes in the innate (DC, monocyte and NK cell) and adaptive (B, CD4^+^ T and CD8^+^ T cell) compartments between children and non-ICU adults during Acute and Convalescent phases. (B) Dotplot showing the number of differentially expressed genes in the innate and adaptive immune cell subclusters between children and non-ICU adults during Acute and Convalescent phases. (C) Expression of interferon response genes by innate immune cells from children, non-ICU and ICU adults during Acute and Convalescence. (D) Expression of interferon response genes by adaptive immune cells from children, non-ICU and ICU adults during Acute and Convalescence.

The most consistently upregulated genes were interferon-induced genes. We therefore generated an interferon gene signature (**Table S3**), and scored each cell subpopulation in children, non-ICU adults and ICU adults (**Fig. 3C** and **3D**). These data confirmed the high expression of interferon-induced genes, particularly in the monocyte subpopulations and novel interferon-activated naïve CD4^+^ and CD8^+^ T cells. Interferon gene signatures were elevated in the acute phase and persisted in the convalescent phase, and was higher in non-ICU adults than children. These data show that SARS-CoV-2 infection leaves a more profound immunological footprint, especially in the adaptive B and T cell compartments, in adults compared to children with the same disease severity.

### Differences between SARS-CoV-2-specific T cells in children and adults

We next analyzed the 158,975 CD4^+^ T cells and 105,273 CD8^+^ T cells where we were able to sequence and reconstruct the TCR. There was similar clonal diversity in the CD4^+^ T cell compartment in children and adults as measured by Shannon Entropy index (**Fig. 4A**). However, there were significant differences in the CD8^+^ T cell compartment where children had the most diverse TCRs followed by non-ICU adults and then ICU adults (**Fig. 4A**). Within each group the diversity did not differ between acute and convalescent timepoints. TCR sequences were annotated using ImmuneCODE (Nolan et al., 2020) and VDJdb (Bagaev et al., 2020), large-scale databases of TCR sequences and binding associations with the addition of SARS-CoV-2 TCR sequences reported in the literature and the SARA-CoV-2-annotated cells mapped on to the T cell clusters in the UMAP for children, non-ICU adults and ICU adults (**Fig. 4B**). The ImmuneCODE database aggregates >135,000 TCR sequences that have been shown with high confidence to react against SARS-CoV-2, while VDJdb collects and curates TCRs of diverse specificity from direct submissions and mining of the literature. These databases do not represent the full landscape of SARS-CoV-2 specificity as they are highly dependent on the peptide pools used to stimulate the T cells. We therefore sought to explore additional SARS-CoV-2 responding T cells within our donors by bulk TRB sequencing pools of proliferated T cells that had been stimulated with recombinant S and RBD proteins. Clonotypes with enriched frequencies after *in vitro* antigen-specific expansion were considered to have putative SARS-CoV-2 specificity (**Table S4**). In total, we annotated significantly more SARS-CoV-2- specific T cells in children (431 of 81,686 in acute and 474 of 84,567 cells in convalescence) than non-ICU (209 of 46,216 in acute and 196 of 49,620 in convalescence) and ICU adults (132 of 17,081 in acute and 102 of 19,292 in convalescence) (p<0.0001, Fisher’s exact test). We matched 746 CD4^+^ and 798 CD8^+^ T cells in our dataset that were potentially reactive against the envelope, surface glycoprotein, membrane glycoprotein, nucleocapsid phosphoprotein, ORF1ab, ORF3a, ORF6, ORF7a, ORF7b, ORF8 and ORF10 of SARS-CoV-2 (**Fig. 4C** and **4D**). Children harbored more SARS-CoV-2-annotated CD4^+^ T cells (mean 76 cells, 74 clones) than non-ICU (mean 49 cells, 49 clones) and ICU adults (48 cells, 40 clones); however, these T cells had similarly diverse TCR repertoires capable of recognizing multiple components of the virus (**Fig. 4C**). Notably, both ICU adults had clonally expanded SARS-CoV-2-annotated CD4^+^ T cells. The differences were more apparent in the CD8^+^ T cell compartment where children had more SARS-CoV-2-specific CD8^+^ T cells (mean of 75 cells, 66 clones in children compared to mean of 41 clones in non-ICU adults and 12.5 in ICU adults) and these included a small number of expanded clonotypes in four of six children (**Fig. 4D**). SARS-CoV-2-annotated CD8^+^ T cell clonal expansions were more marked in several adults, particularly ICU adults where the SARS-CoV-2 annotated repertoire in both acute and convalescent phase was dominated by a few clones.

**Figure 4.**
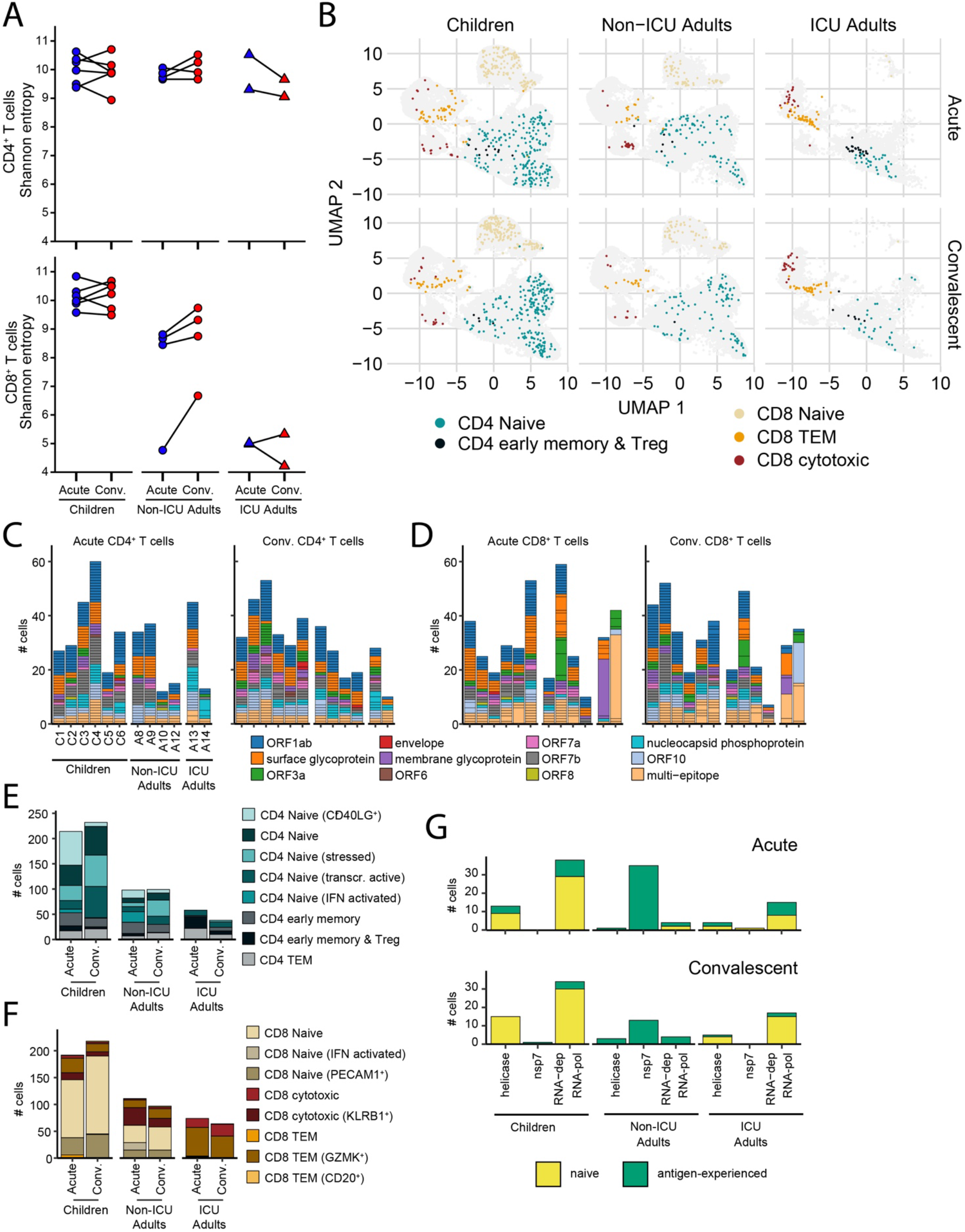
Clonal analysis of SARS-CoV-2-specific T cells. (A) Shannon entropy score for CD4^+^ (top) and CD8^+^ (bottom) T cells in children, non-ICU and ICU adults in Acute and Convalescent phases. (B) Transcriptional state of T cells annotated as SARS-CoV-2-specific in children, non-ICU and ICU adults in Acute (top) and Convalescent (bottom) phases. (C) Epitope specificity of CD4^+^ T cells for different components of SARS-CoV-2 virus in children, non-ICU and ICU adults in Acute (left) and Convalescent (right) phases. Each stack in the stacked barplot represents a single clone. (D) Epitope specificity of CD8^+^ T cells for different components of SARS-CoV-2 virus in children, non-ICU and ICU adults in Acute (left) and Convalescent (right) phases. Each stack in the stacked barplot represents a single clone. (E) Transcriptional state of SARS-CoV-2-annotated of CD4^+^ T cells in children, non-ICU and ICU adults in Acute and Convalescent phases. (F) Transcriptional state of SARS-CoV-2-annotated of CD8^+^ T cells in children, non-ICU and ICU adults in Acute and Convalescent phases. (G) Number of RTC-specific T cells in children, non-ICU and ICU adults in Acute (top) and Convalescent (bottom) phases. Naïve CD4^+^ and CD8^+^ T cells are yellow and antigen-experienced T cells are green.

We aggregated the SARS-CoV-2-annotated CD4^+^ T cells and noted that this compartment in children comprised predominantly naïve cell clusters (naïve, CD40LG^+^, stressed, transcriptionally active and interferon-activated) at both the acute and convalescent timepoints (**Fig. 4E**). Adults, whose repertoire was dominated by virus-specific TEM, early memory and Tregs, had significantly less naïve and more antigen-experienced SARS-CoV-2-annotated CD4^+^ T cells compared to children (351 naïve, 102 antigen-experienced in children; 133 naïve, 64 antigen-experienced in non-ICU; 25 naïve, 71 antigen-experienced in ICU adults; p<0.001, Chi-square). These differences were more pronounced in the CD8^+^ T cell repertoire where children had more naïve cells than adults (329 naïve, 123 antigen-experienced in children; 118 naïve, 90 antigen-experienced in non-ICU; 4 naïve, 134 antigen-experienced in ICU adults; p<0.001, Chi-square) (**Fig. 4F**).

We also analyzed for the expression of TCRs directed against conserved early components of the SARS-CoV-2 replication-transcription complex (RTC) encoded within ORF1ab, including RNA-polymerase cofactor non-structural protein 7 (NSP7), RNA-dependent RNA polymerase (NSP12) and RNA helicase (NSP13), that have been proposed to cross-react with hCoV (Swadling et al., 2021). Children had more RTC-specific T cells and these were predominantly naïve compared to non-ICU adults where they were almost exclusively antigen-experienced and ICU adults where they were a mix of naïve and experienced T cells (**Fig. 4G**). Interestingly, the transition from acute to convalescence was associated with attrition of RTC-specific antigen-experienced T cells in both children and adults. Taken together, these data reveal differences in the SARS-CoV-2-specific T cell compartment in children and adults that may reflect prior antigen exposure to cross-reactive hCoV and virus-induced changes to T cell composition and repertoire.

### Annotation and tracking of T cell clonotypes in children and adults

Analysis of the annotated clonotypes revealed that the majority of T cells were unique and only could be detected at either the acute or convalescent timepoints, but rarely both (**Fig. 5A**). T cell clonotypes that could be tracked longitudinally were more prevalent in the CD8^+^ T cell compartment in ICU adults. The CD4^+^ compartment was almost entirely comprised of unique clonotypes (single cells) with a higher proportion of clonotypes being expanded (> 1 cell) within the CD8^+^ T cell compartment for both the children and adults (**Fig. 5B**). Notably, adults had a higher frequency of expanded clonotypes compared to children, especially the ICU adults. Clonally expanded T cells detected at both acute and convalescent timepoints were considered longitudinal clones.

**Figure 5.**
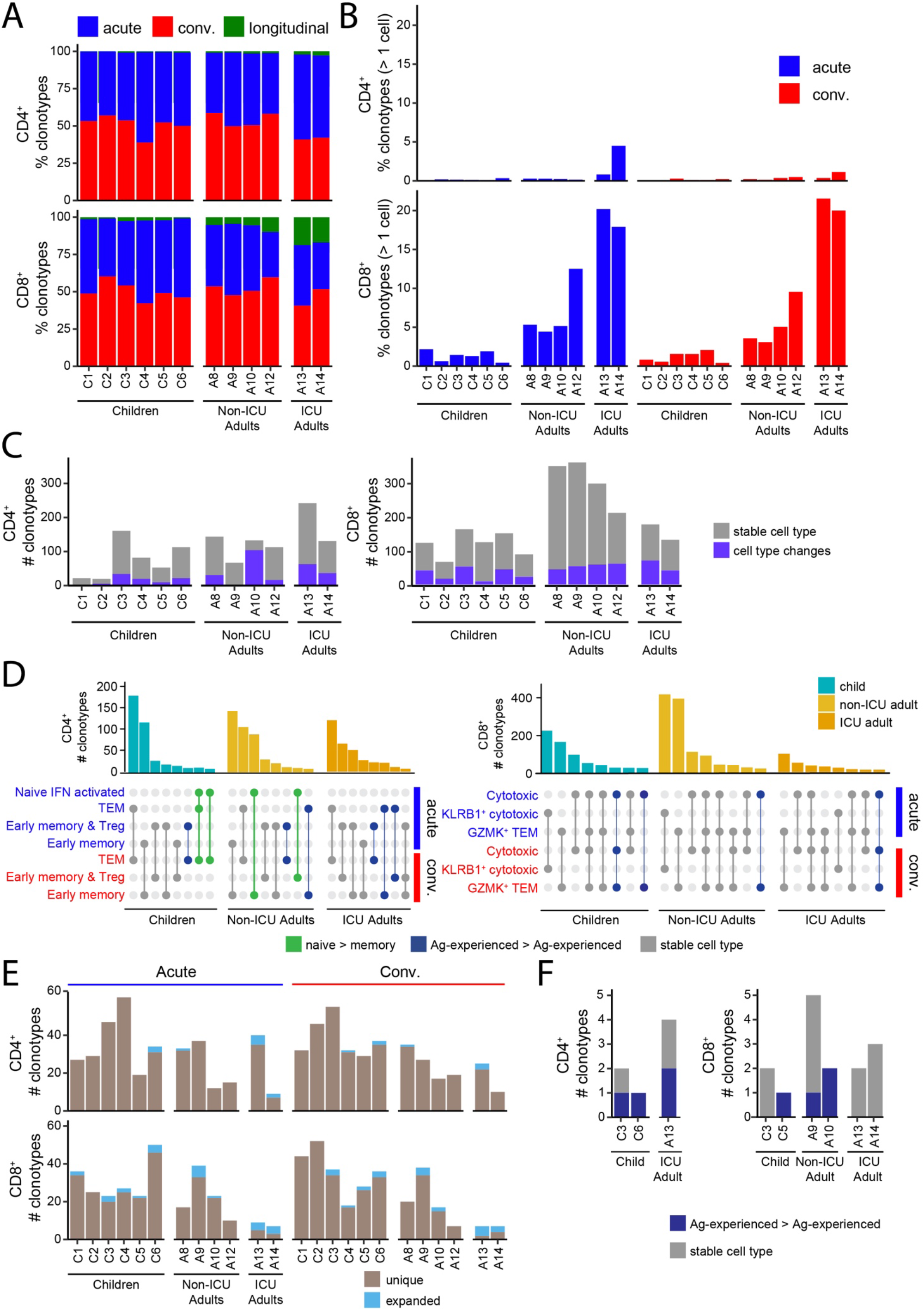
Clonal dynamics of SARS-CoV-2-specific T cells. (A) Percentage of CD4^+^ (upper) and CD8^+^ (lower) clonotypes that are unique at the Acute (red) or Convalescent (blue) phase of infection or present in both (green) for each subject. (B) Number of CD4^+^ (upper) and CD8^+^ (lower) clonotypes that are detected at both the Acute and Convalescent phase of infection (longitudinal clonotypes). (C) Counts for CD4^+^ (left) and CD8^+^ (right) clonotypes coloured by whether their cell type remains the same (grey) or changes (purple) between the Acute and Convalescent phases. (D) Top 8 cell type distributions for CD4^+^ (left) and CD8^+^ (right) longitudinal clonotypes for children, non-ICU adults and ICU adults. The barplots indicate the number of clonotypes with the cell type distribution pattern depicted below each bar where a filled circle indicates that the cell type on the y- axis is present. Distributions are coloured to indicate whether they represent transitions from naïve to antigen-experienced (green), transitions between antigen-experienced compartments (blue) or are the same cell type across the two timepoints (grey). (E) Clonotype counts for SARS-CoV-2-annotated clonotypes for all donors for the CD4^+^ (upper) and CD8^+^ (lower) compartments coloured by whether the clonotype was unique (light brown) or expanded (light blue) at either the Acute (left) or Convalescent (right) phase. (F) Clonotype counts for longitudinal SARS-CoV-2-annotated CD4^+^ (left) and CD8^+^ (right) T cells for the subset of donors that harbor them. Clonotypes are grouped and coloured by whether they have the same cell type at both timepoints (grey) or altered their cell types between Acute and Convalescent phases (dark blue).

A small number of CD4^+^ and CD8^+^ T cell clones were detected in both the acute and convalescent phase in both children and adults making it possible to track their trajectories (**Fig. 5C**). The CD4^+^ T cells tended to be antigen-experienced and unique at the acute timepoint. In contrast, the CD8^+^ T cells were almost exclusively made up of antigen-experienced cells which were a mix of both unique and clonally expanded cells in the acute phase. We tracked the transcriptional state of these longitudinal clones to determine if they had undergone cellular differentiation and found that most clonotypes were stable with few transitions from one T cell state to another (**Fig. 5C**). We identified 27 out of 38,051 (0.07%) naïve CD4^+^ T cells in children that transitioned to TEM and 97 out of 15,022 (0.65%) in non-ICU adults that transitioned to early memory T cells (p<0.0001, Fisher’s exact test) (**Fig. 5D**). Interestingly, child TEM and non-ICU adult CD4^+^ early memory T cells all originated from the novel interferon-activated naïve CD4^+^ T cell pool. These cells were not annotated as SARS-CoV-2-specific in the ImmuneCODE and VDJdb databases but, given their differentiation trajectory, may represent clonotypes responding to the virus. No transitions from naïve to memory were identified in the CD8^+^ T cell clonotypes. Thus, naïve CD4^+^ T cells were significantly more likely to differentiate into memory T cells in adults than children with the same disease severity.

### Longitudinal tracking of SARS-CoV-2-annotated T cells in children and adults

We next examined the trajectories of SARS-CoV-2-annotated T cells. This revealed that in children they were rarely clonally expanded, consisting of 1.39% of acute and 1.30% of convalescent CD4^+^ T clones, and 6.52%, of acute and 4.19% of convalescent CD8^+^ T clones (**Fig. 5E**). Even among the non-ICU adult CD8^+^ compartment, clonal expansions of the SARS-CoV-2-annotated cells were rare (acute 7.87%, convalescent 7.32%) (**Fig. 5E**). The most evidence for SARS-CoV-2-specific T cell clonal expansion was in the ICU adult CD8^+^ compartment where 50% and 57% of the SARS-CoV- 2-annotated clonotypes were expanded at both sampling timepoints, respectively. Only 6 or 7 clonotypes, equating to 0.71% and 1.95% in kids and non-ICU adults, respectively, were found at both timepoints (**Fig. 5F**). Of the expanded clones, 25% of the CD8^+^ clonotypes from the ICU adults were longitudinal cloned present at both timepoints. For ICU adult A13, the 7 days between sampling may make repeated detection on the same clonotypes more likely, but this was also observed for ICU adult A14, who was sampled 149 days apart. These CD8^+^ SARS-CoV-2-specific T cells showed no evidence of phenotypic differentiation (**Fig. 5F**). Taken together, these data suggest that children have a large pool of diverse, polyclonal naïve virus-specific T cells that largely remain intact, whereas the adult repertoire include clonally expanded memory T cells that undergo activation and attrition, possibly via terminal differentiation.

### Functional state of SARS-CoV-2-specific T cells in children and adults

The TCR annotations enabled further analysis of the interferon response gene signature that was differentially expressed at the global level between T cells from children and non-ICU adults (**Fig. 3E**). The interferon response gene signature was enriched in SARS-CoV-2-specific T cells and we were able to detect high scores in cells, such as subpopulations of CD8^+^ T cells, that were not evident at the global level (**Fig. 6A**). Interferon-activated naive CD4^+^ T cells and CD20^+^ CD8^+^ T cells from non-ICU adults had the highest interferon response scores. We were also able to annotate a number of TCR specificities against cytomegalovirus (CMV), Epstein-Barr virus (EBV) and influenza. There were sufficient numbers of CMV-specific T cells (138 CD4^+^ and 271 CD8^+^ clones), but not enough EBV- or influenza-specific T cells, to enable their analysis. No CMV-specific T cell clones were detected in ICU adults. Interestingly, we detected interferon-response gene activation during the acute phase in a small number of CMV-specific cytotoxic KLRB1^+^ cytotoxic and GZMK^+^ TEM (**Fig. 6A**). This suggests that there may be bystander T cell activation, particularly in acute non-ICU adults where the score was higher.

**Figure 6.**
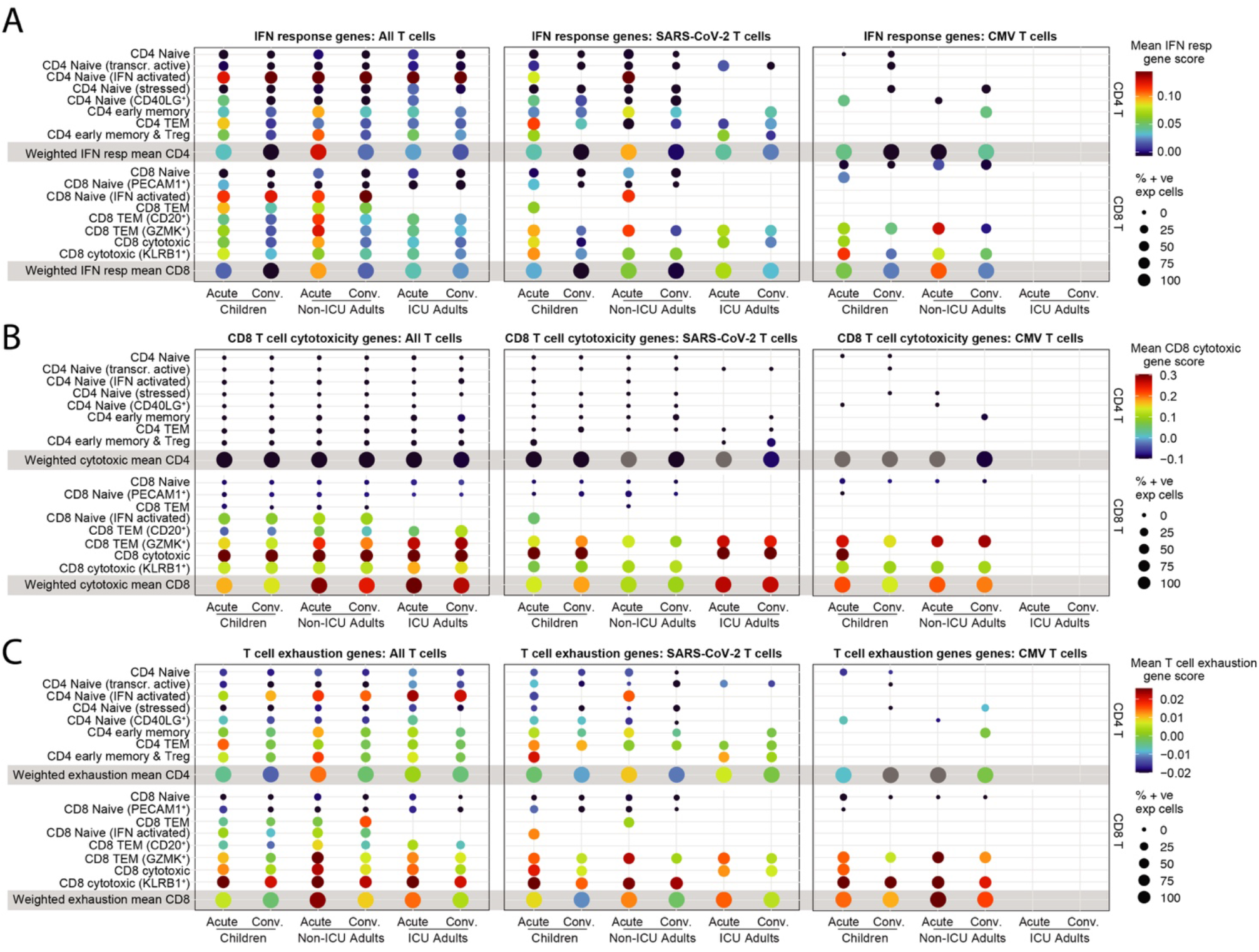
T cell interferon activation, cytotoxicity and exhaustion states. (A) Interferon response gene scores for all T cells (left), SARS-CoV-2-specific T cells (middle) and CMV-specific T cells (right) in children, non-ICU and ICU adults in Acute and Convalescent phases. (B) CD8^+^ T cell cytotoxicity gene scores for all T cells (left), SARS-CoV-2-specific T cells (middle) and CMV-specific T cells (right) in children, non-ICU and ICU adults in Acute and Convalescent phases. (C) T cell exhaustion gene scores for all T cells (left), SARS-CoV-2-specific T cells (middle) and CMV-specific T cells (right) in children, non-ICU and ICU adults in Acute and Convalescent phases.

To further investigate the functional state of SARS-CoV-2-specific T cells, we generated a cytotoxicity gene signature (**Table S4**). Globally, this signature was strongest in CD8^+^ cytotoxic T cells, and stronger in adults, particularly ICU adults, than children, and absent in the CD8^+^ naïve T cell and CD4^+^ T cell populations (**Fig. 6B**). Interestingly, KLRB1^+^ cytotoxic T cells, despite their expression of cytotoxicity genes such as *GZMK, GZMA, NKG7, CST7* and *GNLY* (**Table S2**), did not have a high cytotoxicity score. CD8^+^ GZMK^+^ TEM cells in adults expressed a higher cytotoxicity score than children. Overall, there was evidence for increased cytotoxicity in adults compared to children, and this was most noticeable during acute infection. We also detected upregulation of cytotoxic genes in CMV-specific effector cells in the acute stage in children and in both stages in non-ICU adults.

We next generated a T cell exhaustion signature (**Table S4**). T cell exhaustion scores were significantly higher in CD8^+^ than CD4^+^ T cells where it was predominantly expressed by CD4^+^ interferon-activated naïve T cells (**Fig. 6C**). Expression was highest in ICU adults, followed by non-ICU adults and then children. Expression was higher during acute infection than in convalescence. In the CD8^+^ T cell compartment there was evidence for exhaustion of the GZMK^+^ TEM exhausted memory precursors and cytotoxic T cell subpopulations, particularly the KLRB1^+^ cytotoxic T cells.

Exhaustion scores were higher during acute infection than convalescence and higher in non-ICU adults than children. This pattern was also evident in the SARS-CoV-2- and CMV-annotated T cells, but the smaller number of cells in these groups meant that several cell types were missing. Taken together, these data suggest that SARS-CoV-2-specific T cells in children are less activated, less cytotoxic and less exhausted than their adult counterparts.

### Memory T cell responses to SARS-CoV-2 in children and adults

To determine the functional consequences of these age-specific differences in T cell composition and transcriptional state we performed *in vitro* stimulation with recombinant RBD and S protein to detect antigen-induced upregulation of CD25 and CD134 (OX40) in CD4^+^ T cells (Zaunders et al., 2009). This assay specifically detects CD45RO^+^ memory T cells that are activated after secondary stimulation and not naïve T cells that have not encountered antigen before (Phetsouphanh et al., 2014). This analysis showed that during acute infection, children had variable responses to RBD, which did not significantly change upon recovery (**Fig. 7A**). In contrast, paired samples from non- ICU adults showed a consistent increase in the memory CD4^+^ T cell response to RBD in all patients. A similar pattern was observed in the CD4^+^ T cell response to S protein, with no significant increase in children in contrast to the uniform increase in paired samples from all non-ICU adults (**Fig. 7B**). These differences were reflected in the T cell proliferation assay, which showed significant increase in proliferative responses to RBD (**Fig. 7C**) and S protein (**Fig. 7D**) in adults but not children. There was a moderately positive correlation between the acquisition of memory CD4^+^ T cell responses in the convalescent phase and age to RBD (**Fig. 7E**) and S antigen (**Fig. 7F**). There was also a moderately positive correlation between responses to RBD and the number of SARS-CoV-2-specific memory CD4^+^ T cells (**Fig. 7G**) and a trend towards correlation for responses to S protein (**Fig. 7H**). These data are consistent with the single cell transcriptomic and TCR repertoire analysis and show that natural infection with SARS-CoV-2 induces T cell memory in adults more efficiently than in children.

**Figure 7.**
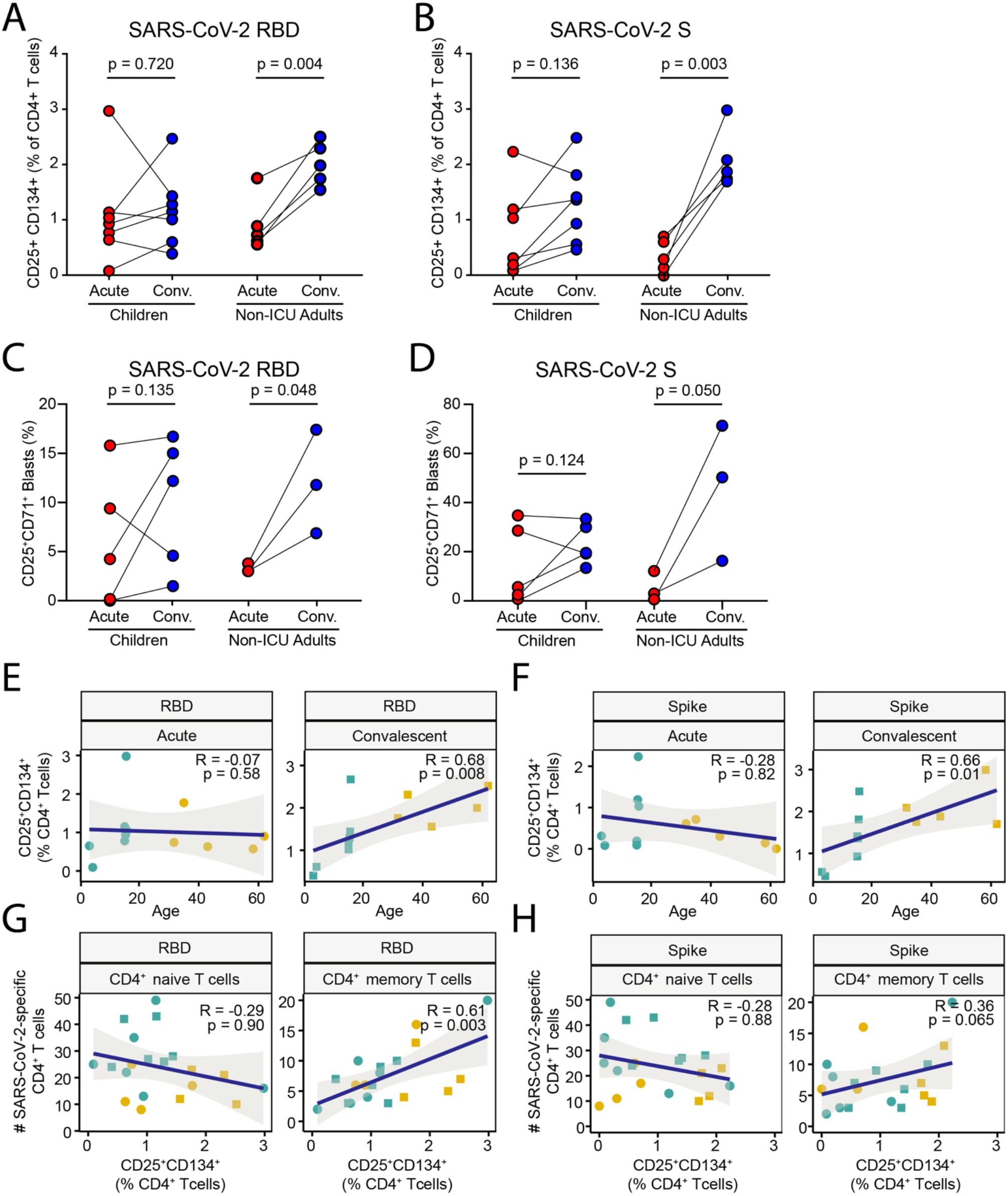
Memory T cell responses to SARS-CoV-2 in children and adults. (A) Frequency of CD25^+^CD134^+^CD4^+^ T cells in cultures of PBMCs stimulated with recombinant SARS-CoV-2 RBD protein from children and non-ICU adults in Acute and Convalescent phases. (B) Frequency of CD25^+^CD134^+^CD4^+^ T cells in cultures of PBMCs stimulated with recombinant SARS-CoV-2 S protein from children and non-ICU adults in Acute and Convalescent phases. (C) CD4^+^ T cell proliferative response in cultures of PBMCs stimulated with recombinant SARS- CoV-2 RBD protein from children and non-ICU adults in Acute and Convalescent phases. (D) CD4^+^ T cell proliferative response in cultures of PBMCs stimulated with recombinant SARS- CoV-2 S protein from children and non-ICU adults in Acute and Convalescent phases. (E) Linear regression of CD25^+^CD134^+^ response to RBD protein by CD4^+^ T cells with age in Acute (left) and Convalescent (right) phases. (F) Linear regression of CD25^+^CD134^+^ response to S protein by CD4^+^ T cells with age in Acute (left) and Convalescent (right) phases. (G) Linear regression of CD25^+^CD134^+^ response to RBD protein by CD4^+^ T cells with number of SARS-CoV-2-specific naïve (left) and memory (right) T cells. (H) Linear regression of CD25^+^CD134^+^ response to S protein by CD4^+^ T cells with number of SARS- CoV-2-specific naïve (left) and memory (right) T cells.

## DISCUSSION

The immune response to SARS-CoV-2 and the immunopathogenesis of severe life-threatening COVID-19 has been the focus of intense investigation in the two years since the first cluster of pneumonia cases were reported in Wuhan, China on the 31^st^ of December, 2019. Important insights have derived from the application of innovative single cell technologies and tissue sampling techniques to deconvolute the local and systemic immune response (Tian et al., 2021). While initial studies have examined adults across the disease severity spectrum, it is only recently that efforts have been directed more towards understanding the immune response of children exposed to SARS-CoV-2. These studies have contributed to a detailed picture in which children are able to rapidly eliminate the virus due to their higher steady state expression of interferon genes and pre-activated innate immune system, especially in the upper respiratory tract (Loske et al., 2021; Yoshida et al., 2021). Similar local innate immune defense mechanisms may operate to protect children from SARS and Middle Eastern Respiratory Syndrome (MERS) to which they are also less susceptible (Rajapakse and Dixit, 2021; Zimmermann and Curtis, 2020). However, this innate resistance to SARS-CoV-2 infection may come at a cost and it is still unclear how the rapid clearance of viral antigens impacts on the adaptive immune response and the generation of immunological memory in children. This is particularly relevant as there is emerging public health concerns over the relative merits and risks of infection- versus vaccine-induced immunity in children. Here, we have concentrated on the systemic immune response in children and adults with the same mild/asymptomatic disease and tracked responses during acute infection and in the convalescent recovery phase to ascertain the factors that may contribute to age-specific differences in COVID-19 severity and its consequences for SARS-CoV-2 immunity. Our longitudinal study avoids any confounding effects from studying heterogeneous patients suffering from varying disease severity.

Our multimodal analysis of the acute and convalescent immune response revealed that COVID-19 leaves a deeper immunological footprint in the adaptive immune system in adults than children. While both children and adults make similar antibody responses against S protein, adults had more circulating activated CD38^+^ HLA-DR^+^ T cells. CD38 and HLA-DR are classical markers of viral infection which may also be induced by bystander activation (Jia et al., 2021; Kim and Shin, 2019) and trogocytosis (Jia et al., 2021). The ICU adults with the highest frequency of activated CD38^+^ HLA-DR^+^ T cells also had elevated serum IL-6 levels. Deconvolution of the circulating immune compartment by high dimensional flow cytometry and single cell RNA sequencing revealed age-specific differences in cellular composition of the NK, B and T cell compartments. However, we also detected widespread upregulation of interferon-induced genes in both the innate (monocytes and NK cells) and adaptive compartment (B, CD4^+^ and CD8^+^ T cells) in adults during acute infection, but this signature was largely limited to the innate compartment in children. This may reflect differences in timing with early production of interferon in children and late interferon production in adults. The importance of interferons in antiviral immunity and resistance to SARS-CoV-2 infection has been well recognized (Lee and Shin, 2020). What was surprising was the fact that interferon gene signatures were largely restricted to the innate immune compartment, suggesting that SARS-CoV-2 infection left only a small immunological footprint in the circulating B and T cells in children. These data are consistent with evidence for reduced breadth of SARS-CoV-2 antibodies in children (Weisberg et al., 2021).

Our analysis also revealed novel subpopulations of naïve T cells, including interferon-activated naïve CD4^+^ and CD8^+^ T cells which were expanded during acute infection and declined in the convalescent recovery phase in children, but which nevertheless were also detectable in healthy adults. Interestingly, naïve T cells exposed to interferon or interferon-induced cytokines, such as IL-15, exhibit signs of activation but do not undergo cell proliferation (Tough et al., 1999). In this regard, it is notable that persistent secretion of type I and type III interferon and activated naïve T cells have been reported in patients with post-COVID-19 syndrome (Phetsouphanh et al., 2022) and it will be interesting to determine if such patients have persistent expansion of interferon-activated naïve CD4^+^ and CD8^+^ T cells. Interferon activation of naïve CD8^+^ T cells have been postulated to enhance their homing, survival, differentiation, antiviral and antibacterial effector functions (Jergovic et al., 2021; Urban et al., 2016). Importantly, clonal tracking revealed that interferon-activated CD4^+^ T cells were the precursors of TEM in children and CD4^+^ early memory T cells in adults. These transitioning expanded T cell clones may represent unannotated SARS-CoV-2-specific T cells. Interferon- activated naïve T cells are a novel cell population and add to the growing recognition from single cell analyses that seemingly homogeneous cell populations, in this case the naïve T cell pool, may be more heterogeneous than previously recognized (Nguyen et al., 2018; Tian et al., 2021). Furthermore, this suggests that traditional nomenclature based on cell surface markers may be inadequate.

We simultaneously sequenced the transcriptome and TCR repertoire of circulating T cells to characterize the SARS-CoV-2-reactive T cell compartment and the impact of COVID-19 on T cell fate and antiviral memory. Children had more diverse TCR repertoires than adults, consistent with their immunological age and previous reports (Naylor et al., 2005; Yoshida et al., 2021). Interestingly, the SARS-CoV-2-specific T cell compartment in children consisted predominantly of naïve T cells with a diverse repertoire capable of recognizing multiple T cell epitopes, including T cells that recognize components of the RTC. RTC-specific memory T cells have recently been proposed to cross-react with hCoV and mediate immunity during abortive SARS-CoV-2 infection in adult health care workers (Swadling et al., 2021). These data suggest that cross-reactive antigen-specific T cells generated by VDJ recombination during ontogeny may still be naïve and not yet selected by antigen experience into the memory pool, particularly in young children who have not been repeatedly exposed to cross-reactive hCoV (Gorse et al., 2020; Pierce et al., 2020; To et al., 2020). In contrast, adults harbored fewer SARS-CoV-2-specific T cells and these included a large number of clonally expanded memory T cells, including to the RTC, that may have been selected by prior infection with hCoV. Such pre-existing cross-reactive T cell memory has been implicated as a risk factor for severe COVID-19 in the elderly (Bacher et al., 2020). On the other hand, recent infection with endemic hCoV have also been associated with less severe COVID-19, possibly by “back-boosting” pre-existing immunity (Sagar et al., 2021).

In addition to the differences in cellular composition, we also detected differences in the transcriptional state of the T cells in children and adults. It has been reported that patients admitted to ICU with severe COVID-19 may have impaired cytotoxicity compared to non-ICU patients as measured by reduced secretion of IL-2 and interferon-*γ* following *in vitro* polyclonal stimulation (Mazzoni et al., 2020). However, our analysis showed that ICU adults had enhanced cytotoxicity scores (particularly in SARS-CoV-2-specific CD8^+^ T cells) compared to non-ICU adults, whereas children had the lowest cytotoxicity scores. This cytotoxicity score encompasses 67 genes that includes, but are not limited to, IL-2 and interferon-*γ*. Increased T cell cytotoxicity in severe COVID- 19 has also been reported by other investigators (Meckiff et al., 2020). Intriguingly, it has also been reported that severely ill COVID-19 patients show features of impaired T cell exhaustion from the single cell RNA sequencing of expanded T cells generated by *in vitro* stimulation with SARS-CoV- 2 peptide pools (Kusnadi et al., 2021). In contrast, our transcriptomic analysis of unstimulated T cells shows a clear hierarchy of exhaustion with children having the least exhausted CD8^+^ T cells followed by non-ICU and then ICU adults. This is consistent with a number of studies showing evidence of T cell exhaustion in patients with severe COVID-19 (De Biasi et al., 2020; Diao et al., 2020; Laing et al., 2020; Zheng et al., 2020a; Zheng et al., 2020b), although it may be difficult to distinguish exhausted from activated cell states (Rha and Shin, 2021; Wherry and Kurachi, 2015). The exhaustion in both non-ICU and ICU adult T cells was accompanied by mitochondrial dysfunction and increased expression of OXPHOS genes, particularly in the two ICU adults. Metabolic dysregulation in patients with severe COVID-19 has been previously described (Siska et al., 2021; Thompson et al., 2021). Taken together, these data suggest that SARS-CoV-2-specific T cells from adult were more activated and terminally differentiated than children despite having the same disease severity.

One puzzling aspect of our data is the fact that many naïve SARS-CoV-2-specific T cells remained suggesting that they either failed to be activated or were transiently activated and returned to a naïve state. When naïve longitudinal clones were detected we only observed recruitment into the memory pool of small numbers of interferon-activated naïve T cells, and this was largely in non-ICU adults. This may reflect sampling differences between timepoints and also biological differences between children and adults. In adults, there is evidence for activation, cytotoxicity and exhaustion of circulating SARS-CoV-2-specific T cells. Under these circumstances, it is possible that hCoV cross-reactive memory and cytotoxic T cells are preferentially recruited at the expense of naïve T cells into the SARS-CoV-2 response where they undergo terminal differentiation leading to clonal attrition and contraction of the SARS-CoV-2-specific memory and effector T cell pool in convalescence. This exclusion of adult naïve T cells from the response by pre-existing cross-reactive memory T cells is consistent with T cell original antigenic sin (Klenerman and Zinkernagel, 1998). Such imprinting may bias T cell responses and promote the immune escape of SARS-CoV-2 and its variants. The description of pre-existing RTC-specific T cells that expand in adult health care workers with abortive SARS-CoV-2 infection (Swadling et al., 2021) argues against this notion. However, we found antigen-experienced RTC-specific T cells decreased in number in both adults and children, consistent with their clonal attrition. In children, we have shown that, apart from the interferon-activated naïve T cells, there is little evidence for activation of circulating T cells. Therefore, it is possible that naïve SARS-CoV-2-specific T cells are sub-optimally activated in children due to the rapid clearance of viral antigens (Loske et al., 2021; Tosif et al., 2020; Yoshida et al., 2021). Rapid viral clearance may also shut down the secretion of interferon and interferon-induced IL-15, cytokines that have been shown to be needed for the generation of CD8^+^ memory T cells (Kolumam et al., 2005; Schluns and Lefrancois, 2003). This has important implications for the development of T cell memory and the protection afforded by infection-induced immunity in children.

Longitudinal analysis detected the generation of robust CD4^+^ T cell memory responses to RBD and S protein in adults but not children by the OX40 assay, which detects memory and not naïve CD4^+^ T cell responses (Phetsouphanh et al., 2014; Zaunders et al., 2009). In children, this result is consistent with the single cell transcriptomic and TCR data showing little evidence for activation of circulating SARS-CoV-2-specific T cells and recruitment of naïve T cells into the memory pool. Adults also did not show evidence of *de novo* naïve T activation or recruitment but, in contrast to the children, had an expanded SARS-CoV-2-specific memory T cell pool. We noted that OX40 responses were positively correlated with age and the number of SARS-CoV-2-specific memory CD4^+^ T cells. Interestingly, single cell analysis of *in vitro* stimulated PBMCs suggest that severe COVID-19 is associated with increased cytotoxic CD4^+^ and decreased regulatory T cell activity (Meckiff et al., 2020). We note that early memory T cells and Tregs dominated the SARS-CoV-2-specific CD4^+^ T cell compartment in the ICU adults. There was also more evidence of acute systemic T cell activation, cytotoxicity and exhaustion in both SARS-CoV-2-specific and non-specific adult T cells which declined in convalescence. Therefore, the development of RBD and S protein memory T cell responses in adults may not only be due to the emergence of new memory T cells from interferon- activated naïve T cells but also recovery of pre-existing memory T cells from exhaustion or active suppression by regulatory T cells.

Several studies from Hong Kong, Australia and the United States have shown impaired humoral and cellular immune responses in children exposed to SARS-CoV-2 (Cohen et al., 2021; Toh et al., 2021; Tosif et al., 2020; Weisberg et al., 2021). This agrees with our data showing impaired generation of T cell memory responses in children compared to adults. It is possible that the rapid efficient elimination of virus by the innate immune system reduces the antigen availability and prolonged cytokine exposure needed to generate long-lived cellular immunity. Under this scenario, since responses are short-lived, children are dependent on their primed innate immune system for continuing protection from reinfection. These data collectively contradict a recent study showing that children may actually generate more robust adaptive immune responses to SARS-CoV-2 than adults (Dowell et al., 2022). In a large cohort of convalescent primary school children and their teachers from England, it was shown that children make higher titres of cross-reactive antibodies and T cell memory responses than adults. However, this involved a heterogeneous study group with an uneven mix of male and female patients who are not hospitalized but may otherwise suffer from varying degrees of disease severity. Another point of difference is that our study examines the response of a naïve patient group to the second wave of COVID-19 in Sydney which occurred after a period of a time when there was no SARS-CoV-2 cases in the community. The study from England began recruiting after the lockdown was lifted in June, 2020 in the setting of widespread community transmission and may include children and adults who have been multiply exposed to SARS-CoV-2.

A critical question arising from all these studies is the risk of reinfection following COVID-19 in children. Our data supports other studies that would suggest that potent innate immunity undermines the adaptive immune response. If correct this suggests that vaccination, for example with of the BNT162b2 Covid-19 vaccine (Walter et al., 2021), will be required to bypass this immune bottleneck in the upper airway in children and allow them to generate long-lasting immunity.

### Limitations of the study and future directions

There are several limitations to our study. The sample size of our cohort is relatively small and this reduces the confidence in our conclusions. However, this is mitigated by the longitudinal study design and the fact that we have focused on the systemic immune response in patients with the same asymptomatic/mild disease severity from the same households. We longitudinally followed patients from the acute stage to convalescence, and complemented our cohort with an additional two patients with severe COVID-19. We also analyzed 433,301 single cells directly *ex vivo*, making it one of the larger single cell datasets to be made available. Another limitation of our study is the short study period and it will be interesting to see the impact of immunity on intercurrent hCoV and SARS-CoV- 2 reinfection rates with long term follow-up. Accordingly, large-scale prospective studies involving longitudinal study of homogeneous patient groups with the same disease severity will be needed to determine the relative merits and risks of infection- vs vaccine-induced immunity against SARS- CoV-2. Due to technical reasons and the volume of blood sample available from children, we were only able to test memory CD4^+^ T cell responses to RBD and S protein. In addition, the identification of SARS-CoV-2-specific T cells in our study was dependent on the annotation of their TCR in ImmuneCODE and VDJdb databases, which are almost certainly incomplete. Another limitation of our study is the absence of pre-exposure blood samples and viral loads to determine the kinetics of viral clearance, interferon response and baseline cross-reactive immunity to hCoV. Future studies involving more innovative technologies with smaller sample requirements, including nasal sampling of local immune responses, may provide a more complete picture of the dynamic clonal landscape of both local and systemic T cell responses to SARS-CoV-2 and hCOV.

## Acknowledgments

We thank the patients and their families. We thank Miles Davenport, Tony Basten and Robert Brink for critical discussions and comments on the manuscript. We thank Eric Lim, Hira Saeed and staff in the Garvan-Weizmann Centre for Cellular Genomics for technical support. Drawings were created with BioRender.com.

## Author contributions

P.N.B. is the coordinating principal investigator for clinical site. R.N., P.S.H., P.N.B. and T.G.P. conceived and designed the study. R.N., P.S.H., P.N.B., A.H.-J., A.B., B.T., N.W. and D.C. recruited patients and collected patient data. D.R.C. leads the biospecimen research services. L.Z., A.Y., C.L.L., T.V. and R.B. collected and processed patient samples. R.B. managed the project. W.H.K. and T.G.P. designed experiments. W.H.K. performed single cell transcriptome and repertoire sequencing. M.S. performed bulk TCR sequencing. W.H.K., K.J., J.A.-H., S.Y., J.E.P., W.K. and T.G.P. analyzed the sequencing data. K.J. analyzed the TCR repertoire. C.P. and J.J.Z. performed *in vitro* T cell stimulation. S.R.-D. and E.K.D. performed and analyzed the flow cytometry and cytokine bead array. F.L., V.M., F.X.Z.L and F.B. measured anti-S antibodies. R.R. and D.C. generated the recombinant RBD and S proteins. P.I.C., A.K.D., C.G.G., J.E.P., R.N., P.S.H., P.N.B. and T.G.P. provided supervision. W.H.K., K.J., J.D.S., P.I.C., A.K.D., R.N., P.S.H., E.K.D., P.N.B. and T.G.P. wrote the manuscript.

## Supplementary Figure legends

**Figure S1.**
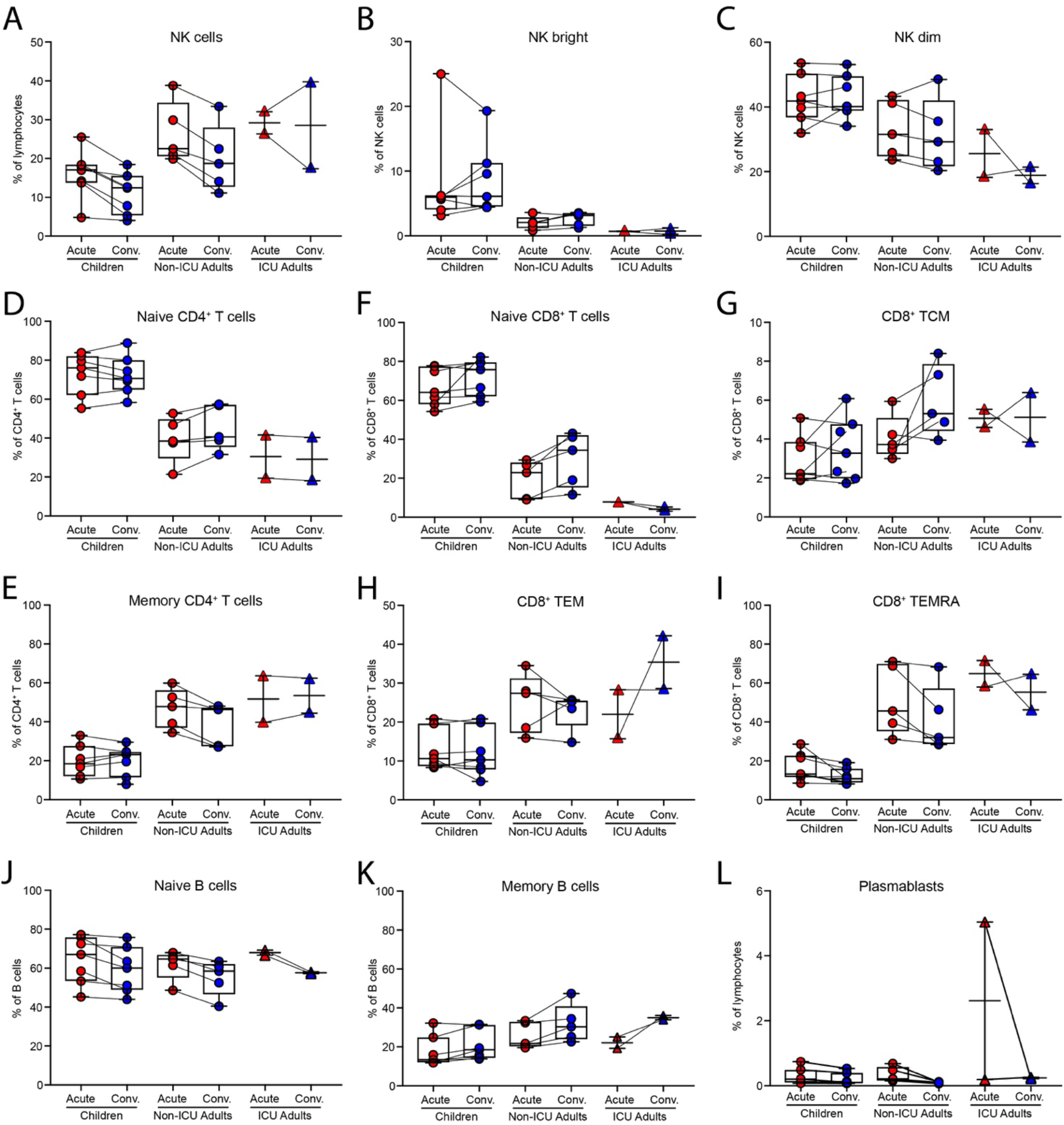
Flow cytometric analysis of PBMCs in children and adults with COVID-19. (A) NK cells. (B) NK bright cells. (C) NK dim cells. (D) Naïve CD4^+^ T cells. (E) Memory CD4^+^ T cells. (F) Naïve CD8^+^ T cells. (G) CD8^+^ central memory T cells (TCM). (H) CD8^+^ effector memory T cells (TEM). (I) CD8^+^ terminally differentiated effector memory T cells (TEMRA). (J) Naïve B cells. (K) Memory B cells. (L) Plasmablasts.

**Figure S2.**
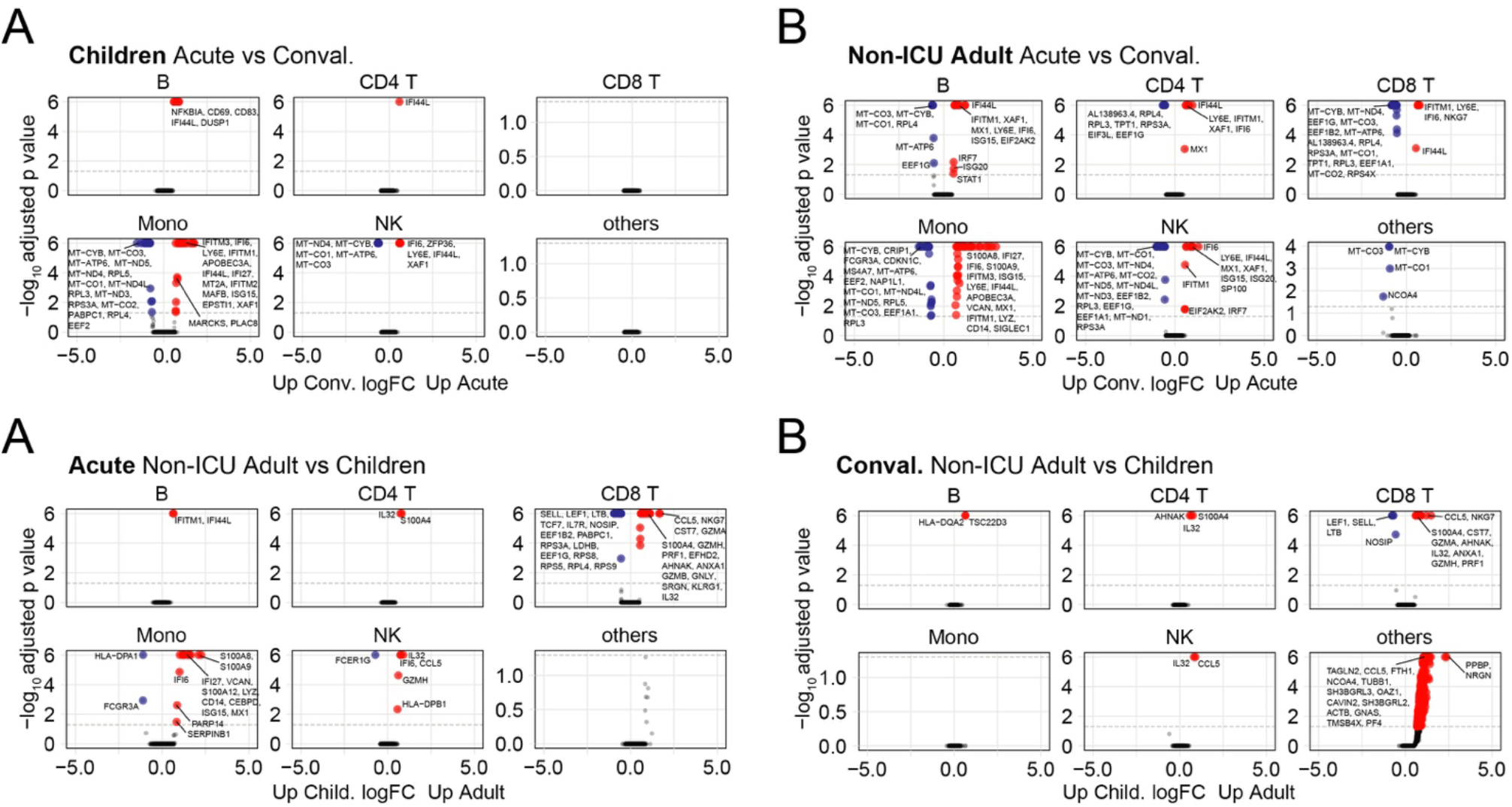
Volcano plots showing top 50 differentially expressed genes. (A) Between acute and convalescent samples in children. (B) Between acute and convalescent samples in non-ICU adults. (C) Between non-ICU adults and children in the acute phase. (D) Between non-ICU adults and children in the convalescent phase.

## METHODS

### EXPERIMENTAL MODELS AND SUBJECT DETAILS

#### Mild/asymptomatic COVID-19 children and adult family members

Participants were recruited under the Immunophenotyping of COVID-19 and other Severe Acute Respiratory Infections Study (2020/PID00920) approved by the Sydney Children’s Hospital Network (SCHN) Human Research Ethics Committee (2020/ETH00837). Potential child participants were identified through admission to the Children’s Hospital at Westmead following a positive SARS- CoV-2 reverse transcriptase polymerase chain reaction (RT-PCR). Families were approached and both children and positive household adults were consented to longitudinal blood sampling. Blood used in these analyses were collected in lithium heparin and serum-separation tubes (Becton Dickinson, USA), and collected both acutely (within 14 days of first positive RT-PCR) and approximately one-month post infection. All patients were ambulant with either mild symptoms, consisting of a cough or sneeze, or asymptomatic (WHO Clinical Progress Scale of 1 out of 10). All consented samples used in these analyses were collected between 23^rd^ July, 2020 and 24^th^ October, 2020.

#### Severe COVID-19 patients

Participants were recruited under the South-East Queensland COVID Consortium study approved by the Gold Coast Hospital and Health Service Human Research Ethics Committee (HREC/2020/QGC/63082). Patient recruitment was undertaken by research team members at the Gold Coast University Hospital. A waiver of informed consent, with the opportunity to opt-out was used for this study. Both patients were elderly males with severe COVID-19 who were intubated and ventilated ICU (WHO Clinical Progress Scale of 7 out of 10). Acute samples were taken on the 14^th^ April 2020. A13 had second sample 8 days later on the 22^nd^ April, 2020 and A14 had convalescent sample taken on the 10^th^ September, 2020.

### METHOD DETAILS

#### PBMC processing and cryopreservation

PBMCs were diluted 1:1 with PBS and the mix overlaid in a 2:1 ratio with Ficoll-Paque in 50ml Falcon tube. Cells were centrifuged at 1700 rpm for 30 minutes with brake off at room temperature (18-21°C). Mononuclear cells at the interface were collected and transferred to a new Falcon tube and washed *×*2 with ice-cold PBS/0.5% FBS by centrifugation at 1300 rpm for 8 minutes with brake on. Cells were resuspended in 5ml culture media (RPMI/1% HEPES) for counting with a haemocytometer and Trypan blue. Cells were then resuspended in 500μl culture media at a concentration of 2 *×* 10^7^/ml and 500μl freezing media (200ml culture media, 200ml FBS and 100ml DMSO) and cryovials transferred to the -80°C freezer in a Styrofoam box for 24-48 hours before storage in liquid nitrogen. Frozen PBMCs were thawed and centrifuged at 400g for 5 minutes. Samples were washed with twice with 4% FBS/PBS and stained with DAPI (0.1 μg/ml) for 5 minutes at room temperature prior sorting of live DAPI^—^ cells on the AriaIII (BD Biosciences)

#### Serum antibody testing

Serum was prepared from clotted tubes by centrifugation at 1000g for 10 minutes. IgG titres to the SARS-CoV-2 Spike protein were measured by our previously published high sensitivity flow cytometry cell-based assay (Tea et al., 2021) which has been modeled on autoantibody detection test used in clinical diagnostic testing of neuroimmunological disorders (Lopez et al., 2022; Tea et al., 2019). HEK293 cells were transfected to express early-clade SARS-CoV-2 Spike antigens. Diluted serum (1:80) was added to live Spike-expressing cells. Codon-optimised, wild-type SARS-CoV-2 strain Wuhan Spike protein ORF with 18 amino acids deleted from the cytoplasmic tail was cloned within the MCS of a lentiviral expression vector, pLVX-IRES-ZsGreen1, using EcoRI and XbaI restriction sites, resulting in pSpike-IRES-ZsGreen vector. All synthetic gene fragments were ordered through IDT. Cells were then incubated with Alexa Fluor 647-conjugated anti-human IgG (H+L) (ThermoFisher Scientific). Cell events were acquired on LSRII flow cytometer (BD Biosciences, USA), and median fluorescence intensity (MFI), a proxy of antibody titres was analysed. The threshold for a positive result was determined if the delta MFI (rMFI = MFI transfected cells – MFI untransfected cells) was above the positive threshold (mean rMFI + 4SD of 24 pre-pandemic age- matched controls) in at least two of three quality-controlled experiments. The sensitivity of the assay was superior to several commercial assays at 98% (95% CI: 92-99%) (Tea et al., 2021). Data were analysed using FlowJo 10.4.1 (TreeStar, USA), Excel (Microsoft, USA) and GraphPad Prism (GraphPad Software, USA).

#### Serum cytokine bead array

We used the BD Cytometric Bead Array (CBA) kit to measure serum cytokines. Cytokine standards were made up in assay diluent as per the manufacturer’s instructions. Capture bead mix was made up in Capture Bead Diluent for Serum/Plasma to a final concentration of 25 µL/test. Serum samples were diluted 1:1 in assay diluent. To each well 25µL of beads was added, followed by 25µL of standard or 25µL of diluted serum sample. Samples were incubated with beads in the dark for 1 hour at room temp. Detection master mix was prepared as per the manufacturer’s instructions and 25µL added to each sample and then samples incubated for a further 2 hours in the dark at room temperature. Beads were then washed twice in wash buffer and then acquired on BD FACS Canto II flow cytometer (BD Pharmingen). Analysis was performed on FCAP array software v 3.0 (BD Biosciences). Cytokine concentrations were imported into R (version 4.1.2), log transformed, and visualized with ggplot2 (version 3.3.5).

#### Flow cytometry

Thawed cells were resuspended in 2% FBS/PBS and plated in a 96-well V-bottom plate. Antibody cocktails were prepared in FACS buffer (0.1% BSA/0.1% sodium azide/PBS). Cells were pelleted by centrifugation at 490g for 5 minutes at 4°C. Cells were then stained with 50µL of Zombie UV Fixable viability dye (diluted 1/500 in PBS) for 20 minutes on ice in the dark. Cells were washed 3 times with FACS buffer. Cells were then incubated with 50µL of blocking agents (normal mouse serum 1/20, Fc block 1/10) for 15 minutes on ice. Antibody cocktails were prepared in FACS Buffer and 50µL added to each sample and then incubated for 30 minutes on ice in the dark. Cells were washed 3 times in FACS buffer and then fixed by resuspending in 150µL of 1% formaldehyde for 20 minutes at room temperature. Cells were then washed and resuspended in FACS buffer and run on FACSymphony (BD Pharmingen). Samples were analysed using FlowJo software (Tree Star).

Intracellular staining for MX-1 was performed for 1 *×* 10^5^ thawed PBMC using the Transcription Factor Buffer Set (BD Biosciences) according to the manufacturer’s directions. Permeabilized cells were stained with CD3-PerCP-Cy5.5, CD4-BUV395, CD8-BUV805, CD45RA-BUV737, CD27-APC-R700 (BD Biosciences) and 1 µg MX-1-AF647 (Abcam) according to manufacturer’s directions and analysed on a 5-laser Fortessa X20 (BD Biosciences) as previously described (Zaunders et al., 2020).

#### Recombinant RBD and S protein

Expression plasmids encoding His-tagged SARS-CoV-2 RBD (residues 319 to 541 of SARS-CoV-2 S protein) or S protein (with a C-terminal trimerization domain) were cloned into pCEP4 vector and transfected into Expi293F (ThermoFisher Scientific) and the proteins expressed for 7 days at 37°C (Rouet et al., 2021). The proteins were captured from the clarified cell culture using TALON resin (ThermoFisher Scientific) and eluted with imidazole. The full trimeric S protein was further purified by size exclusion chromatography (Superose 6 resin) to remove dissociated S1 and S2 domains. The protein purity was assessed by visualization on SDS-PAGE gel.

#### OX40 antigen-specific memory T cell assay

Antigen-specific CD4 T-cells responding to recall antigens were measured in cultures of 300,000 PBMC in 200 µl/well of a 96-well plate, in Iscove’s Modified Dulbecco’s Medium (IMDM; Thermofisher, Waltham, MA, USA) containing 10% human serum (Wayne Dyer, Australian Red Cross Lifeblood, Sydney, Australia), and incubated for 44-48 hr incubation, in a 5% CO2 incubator, as previously described (Zaunders et al., 2009). Separate cultures were incubated with different antigens including: (i) culture medium only negative control well; (ii) anti-CD3/anti-CD28/anti-CD2 T cell activator (1/100 dilution) polyclonal positive control well; (iii) 5 µg/ml recombinant SARS-CoV-2 S trimer; and (iii) 5 µg/ml recombinant SARS-CoV-2 RBD. 100 µl of PBMC from the respective cultures were stained with CD3-PerCP-Cy5.5, CD4-FITC, CD25-APC, and CD134-PE (BD Biosciences, San Jose, CA, USA), and live/dead fixable NIR dead cell stain kit reagent according to manufacturer’s directions and analysed on a 5-laser Fortessa X20 (BD Biosciences) as previously described (Zaunders et al., 2020). Antigen-specific CD4^+^ T cells were gated and expressed as CD25^+^CD134^+^ % of CD3^+^CD4^+^ live T cells as previously described (Zaunders et al., 2009). Cultures were classified as positive for antigen-specific CD4^+^ T cells if the CD25^+^CD134^+^ % of CD4^+^ CD3^+^ T cells was ≥ 0.2% (Hsu et al., 2012).

#### Antigen-specific T cell proliferation and RNA extraction

Antigen-specific T cell proliferation was measured in cultures of PBMC incubated with different controls and antigens in separate wells as above for the OX40 assays, except cells were incubated for 7 days. 100µl of PBMC were then stained with CD3-PerCP-Cy5.5, CD4-FITC, CD25-APC, and CD71-BV650 according to manufacturer’s directions and analysed on a 5-laser Fortessa X20 (BD Biosciences) as previously described (Zaunders et al., 2020). Antigen-specific proliferating CD4^+^ T cells were gated as % of Forward Scatter high and CD25^+^CD71^+^CD4^+^CD3^+^ T cells. Proliferating cells remaining in the cultures were further expanded by incubating for a further 7 days with 20 IU/mL IL-2 (Roche Life Science Products). After expansion, cells from each well were used for RNA extraction, using the Maxwell RSC SimplyRNA Tissue kit and the Maxwell RSC automated extraction system (Promega, Madison, WI) as previously described (Suzuki et al., 2021).

#### Single cell RNA transcriptome and TCR repertoire sequencing

Single cell transcriptomic libraries were generated using the 5’v2 Gene expression and immune profiling kit (10x Genomics). Viable PBMCs were sorted into 2% FBS/PBS and cell counts were performed using a haemocytometer. Up to 40,000 cells were loaded into each lane of Chromium Next GEM Chip K Single Cell Kit (10x Genomics) to achieve a recovery cell number of approximately 20,000 cells. Subsequent cDNA and TCR libraries were generated according to manufacturer’s instructions. Generated libraries were sequenced on the NovaSeq S4 flow cell (Illumina) at Read 1 = 28, i7 index = 10, i5 index = 10 and Read 2: 90 cycles according to manufacturer’s instructions.

#### Transcriptomic analysis

##### Pre-processing of raw sequencing files

Single-cell sequencing data was demultiplexed, aligned and quantified using Cell Ranger (10x Genomics) against the human reference genome (10x Genomics, July 7, 2020 release) with default parameters.

Filtering and quality control was performed using Seurat (Stuart et al., 2019) on raw data containing 522,926 cells where 433,301 cells were retained satisfying thresholds of both <10% mitochondria content and number of genes between 200 and 5000. ‘SCTransform’ was used for normalization with regression of batch, gender and cell mitochrondria content covariates (Hafemeister and Satija, 2019).

##### Annotation of cell identities

Cell annotation was performed using ‘reference-based mapping’ pipeline implemented in Azimuth algorithm in Seurat (Hao et al., 2021). Azimuth annotated T cells were re-clustered and T cell sub- populations were manually annotated based on UMAP clustering and markers defined by ‘FindAllMarkers’ function in Seurat.

##### Differential gene expression analysis

Raw counts from defined cell populations were normalised using scran / scater (Lun et al., 2016; McCarthy et al., 2017) and differential gene expression analysis was performed using Limma voom (Law et al., 2014) with regression of Batch and Gender covariates. DGE analysis was not performed on dendritic cells (all sub populations) and CD8 IFN activated Naïve cell populations due to low cell numbers (<100 cells) sampled in the dataset.

##### Gene signature scores

Gene signature scores (**Table S4**) was generated using ‘AddModuleScore’ function in Seurat. Cell sub-populations with less than 5 cells within sample groups were excluded from the analysis. To compare the gene signatures across different sub-populations within sample groups, gene scores were weighted based on the proportion of positive expressing cells within the sub-population.

The interferon gene signature was generated from aggregating unique interferon response genes sourced from (Hadjadj et al., 2020; Kim et al., 2021; Lee et al., 2020; Szabo et al., 2019). T cell exhaustion signature was sourced from (Utzschneider et al., 2020). CD8^+^ cytotoxic T cell signature was sourced from (Szabo et al., 2019).

#### Analysis of the OneK1K cohort

OneK1K Cohort Study was established to investigate the effects of genetic variation on gene expression at single cell resolution. Original cohort includes more than 1000 individuals recruited from the Royal Hobart Hospital, Hobart Eye Surgeons as well as from the retirement villages within Hobart, Australia prior to the COVID-19 pandemic. We have selected 26 age- and sex- matched individuals from this cohort to compare cell type proportions with the COVID-19 patients. The study was approved by the Tasmanian Health and Medical Human Research Ethics Committee (H0012902). Informed consent was obtained from all participants.

Peripheral blood samples were collected into vacutainer tubes containing either FICOLL™ and sodium heparin (8mL CPT™; BD Australia, North Ryde, NSW; 362753) or K2EDTA (10mL; BD Australia, North Ryde, NSW; Catalogue: 366643). During single cell library preparation equal numbers of live cells were combined for 12-14 samples per pool. Pooled single cell suspensions partitioned and barcoded using the 10X Genomics Chromium Controller and the Single Cell 3’ Library and Gel Bead Kit version 2 (PN-120237). The pooled cells were super-loaded onto the Chromium Single Cell Chip A (PN-120236) to target 20,000 cells per pool. Libraries for all samples were multiplexed and sequenced across five 2x150 cycle flow cells on an Illumina NovaSeq 6000. The Cell Ranger Single Cell Software Suite (version 2.2.0) was used to process data produced by the Illumina NovaSeq 6000 sequencer into transcript count tables. Raw base calls from multiple flow cells were demultiplexed into separate pools of samples. Reads from each pool were then mapped to the GRCh38 genome using STAR (Dobin et al., 2013). Cells for each individual were identified using the Demuxlet computational tool (Kang et al., 2018). The most likely individual for each droplet was determined using the genotype posterior probability estimate from imputation of 265,053 exonic SNPs (R2>0.3 and MAF>0.05). In all approaches, **α** was set to 0.5, assuming a 50/50 ratio and other parameters were kept as default. Droplets which were identified as doublets by both Demuxlet and Scrublet (Wolock et al., 2019) were removed from the dataset.

We used our COVID-19 dataset as a reference to guide the cell type classification of the OneK1K cohort using the Symphony approach (Kang et al., 2021). First, we selected the top 5000 highly variable genes conditioned by batch information and normalized the data using factor normalization and logarithmic transformation as implemented in Seurat. Next, we centered and standardized the normalized gene expression data for the highly variable genes and stored the means and standard deviations for each gene across all cells. We performed singular value decomposition on the scaled data. We applied harmony to align the gene expression embeddings by batch using a theta value of 2 and performing 100 clustering iterations and a maximum of 20 rounds of harmony clustering and correction. We applied the same normalization strategy for the OneK1K data and projected the data onto the reference by scaling the OneK1K data using the reference means and standard deviation and aligning the data using Symphony. Finally, we assign the cell type labels to the OneK1K cells using a lazy K-nearest neighbor classifier with k = 5.

#### TCR repertoire analysis

Following processing with 10X Genomics cellranger vdj (v6.1.2) using the human reference the resulting VDJ contigs were post-processed using stand-alone IgBLAST (v1.14) (Ye et al., 2013) to generate further alignment details. Where a single barcode was associated with more than one chain for either the TRB or TRA loci the VDJ with the highest UMI count was retained. Clonal lineages were defined by IgBLAST called V, J and CDR3 amino acid sequences for both TRA and TRB, if available, or by a single chain if paired chains were not available. Expanded clonotypes were defined within each sample (subject and time point) as those observed across 2 or more cells, while longitudinal clonotypes were those from a subject that were observed at both the acute and convalescent timepoints.

TRBs of reported specificities were collected from immuneCODE Multiplex Identification of T cell Receptor Antigen specificity (MIRA) release 002.2 (Nolan et al., 2020), and VDJdb v2021-09-05 (Bagaev et al., 2020). Additional TRBs were added from (Low et al., 2021), (Lineburg et al., 2021) and (Francis et al., 2022). TRBs were formatted to consistent format, where ambiguous TRBVs reported as a separate entry were created for each TRB.

To account for the private SARS-CoV-2 responses that may not be captured in the public databases, bulk TRB sequencing was undertaken following proliferation of the outputs of the OX40 assays. The bulk sequencing assay was adapted from (Shugay et al., 2014). RNA was reverse transcribed to cDNA that incorporated a 10bp universal molecular identifier using a modification of the SmartSeq2 protocol (Picelli et al., 2014) described in (Massey et al, 2020).

TRBs of reported or inferred specificity were mapped to 10x VDJs by matching of TRB clonotype labels. Where the same TRB was reported to bind multiple epitopes, all epitopes were associated with the TRB clonotype. SARS-CoV-2-annotated clonotypes were defined as any clonotype matching a SARS-CoV-2 reported VDJ regardless of poly-specificity or those observed in the bulk repertoire sequencing from the proliferation assay at an enrichment of at least 64-fold above baseline.

TCR clonotype and annotation data were merged with 10x GEX via cell barcodes. Repertoire metrics were summarised in RStudio (v1.4.1106, RStudio Team (2021). RStudio: Integrated Development Environment for R. RStudio, PBC, Boston, MA URL http://www.rstudio.com) using tidyverse package (Wickham et al., 2019). Shannon entropy was calculated for CD4^+^ and CD8^+^ T cell compartments for each subject to explore the diversity (Shannon, 1948). Clonotype distribution across cell types and time points was explored using Upset plots (Lex et al., 2014) as implemented by the ComplexHeatmap package (Gu et al., 2016).

#### Statistics

Statistical analysis was performed using Prism software (GraphPad) or in R. We used unpaired Student’s t-test to compare between 2 groups and paired Student’s t-tests to compare longitudinal differences within the same individuals. We used the one-way ANOVA with Tukey’s correction for comparisons between multiple groups. We used Fisher’s exact test for 2*×*2 contingency tables and Chi-square for 2*×*3 contingency tablesw. Correlation between variables was measured by Pearson’s correlation coefficient with a one-sided Student’s t-test.

